# Phenotypic Screening Identifies Flunarizine as an Inhibitor of Radiotherapy-Induced Astrocyte Reactivity with Therapeutic Potential in Glioblastoma

**DOI:** 10.1101/2025.07.12.664538

**Authors:** Pauline Jeannot, Rebecca Rosberg, David Lindgren, John C. Dawson, Enrico Pracucci, Cornelia Börjesson-Freitag, Julia Martinez, Vasiliki Pantazopoulou, Maria Malmberg, Karolina I. Smolag, Dimitra Manou, Richard J.R. Elliott, Crister Ceberg, Tracy J. Berg, Henrik Ahlenius, Neil O. Carragher, Alexander Pietras

## Abstract

Radiotherapy is part of the standard-of-care for glioblastoma, yet tumors invariably recur as incurable lesions post-treatment. Recent studies suggest that radiation-induced astrocyte reactivity fosters a tumor-supportive environment, however effective strategies targeting reactive astrocyte phenotypes are lacking. Using a novel image-based assay, we screened over 1,700 small molecule compounds, identifying 29 that inhibit radiation-induced astrocyte reactivity in human astrocytes. Among these, Flunarizine, a calcium-entry blocker approved for migraine treatment, significantly reduced astrocyte reactivity in vitro and in vivo. In a genetically engineered glioblastoma mouse model, combining Flunarizine with radiotherapy markedly improved survival without affecting unirradiated controls, indicating specificity for a radiation-induced phenotype. Mechanistically, Flunarizine inhibited radiation-induced fibrosis in vivo and directly suppressed astrocytic TGF-beta activation in vitro. Notably, Flunarizine treatment had no direct effect on primary glioblastoma cells, emphasizing its microenvironmental specificity. In conclusion, we identified Flunarizine as a promising repurposed compound capable of effectively mitigating radiation-induced astrocyte reactivity and delaying glioblastoma recurrence. This approach offers a viable therapeutic strategy to enhance current glioblastoma treatments.

**Graphical abstract:** 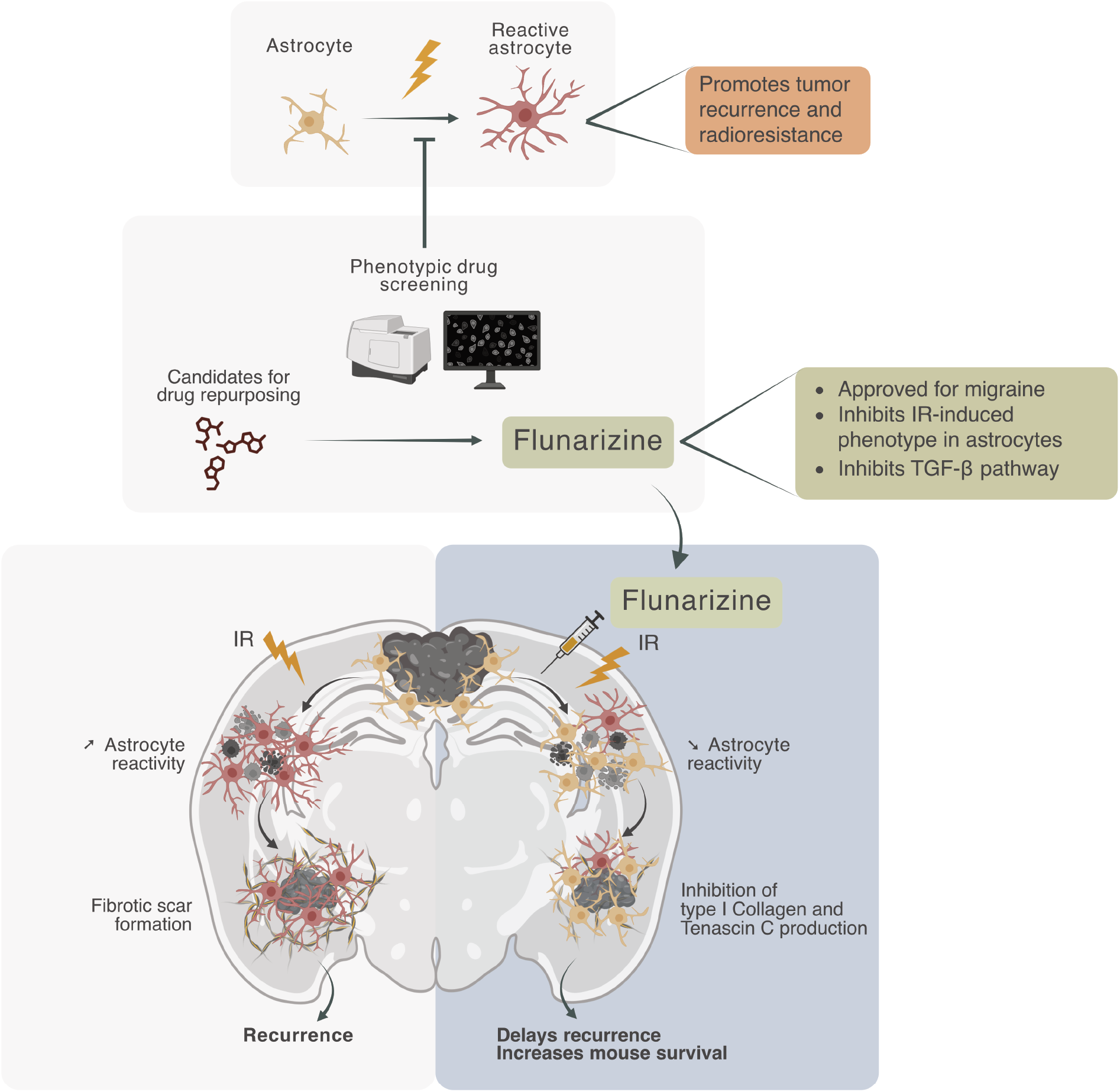

## Introduction

Glioblastomas (GBM) invariably recur as incurable lesions following treatment with the standard of care, which includes chemotherapy, surgery, and radiotherapy (RT) (1). These tumors are highly heterogeneous, exhibiting a range of distinct cell states among tumor cells within the same tumor (2), as well as diverse microenvironmental components, including various glial, immune, and vascular cell types that interact both with each other and with the tumor cells (3). During the course of treatment, various cells of the tumor microenvironment (TME) respond to and adapt as a result of RT. Mounting evidence indicates that RT alters the TME to cause the formation of tumor-supportive conditions in the recurrent tumor (4). Despite an increased understanding of the biological processes underlying such tumor-promoting consequences of RT, few therapeutic strategies have been developed to target microenvironmental stress responses specifically.

Astrocytes represent one of the most prominent cell types in the brain and react to various stresses in a process called reactive astrogliosis. As astrocytes become reactive, they modulate inflammatory processes, remodel the extracellular matrix, and isolate damaged tissue, resulting in protection and repair of neural tissue (5, 6). Sustained or excessive reactive astrogliosis can, however, contribute to pathology including in primary and secondary brain tumors, where reactive astrocytes have been described as tumor-supportive elements of the TME (6, 7). More specifically, tumor-associated astrocytes create an immunosuppressive microenvironment (8, 9), support tumor metabolism (10, 11) and promote brain metastasis (12, 13). We have previously demonstrated that astrocytes, when reactive in response to RT, secrete extracellular matrix components that activate integrin/Src signaling on neighboring GBM cells, and that targeting these interactions can enhance the tumor cell response to RT (14). One clinical study has shown promising results in reducing brain metastasis growth by targeting reactive astrocytes, suggesting that targeting these stress responses may have therapeutic potential also in primary brain tumors (12). These studies further support the pursuit of reactive astrocytes as therapeutic targets in malignant brain tumors.

While reactive astrocytes have been identified as potential therapeutic targets, astrocyte reactivity encompasses a spectrum of context-dependent phenotypes (5) and there are no consensus drugs to target these states universally. Here, we leveraged reactivity-associated morphological changes induced by ionizing radiation (IR) to develop a high-content image-based approach to monitor IR-induced astrocyte reactivity, using both primary human and iPSC-derived astrocytes. Based on this approach, we performed phenotypic screens to evaluate the ability of more than 1,700 experimental and approved small molecule compounds to inhibit IR-induced astrocyte reactivity, identifying 29 compounds for further preclinical evaluation. Among these, the anti-migraine compound Flunarizine consistently reduced IR-induced astrocyte reactivity across model systems, both *in vitro* and *in vivo*. Treatment with Flunarizine prolonged survival and reduced the formation of fibrosis-like scars in a preclinical murine model of GBM. Together, our findings identify Flunarizine as a modulator of astrocyte reactivity and a promising therapeutic approach for GBM.

## Material and methods

### Sex as a biological variable

Animal, cell line, and clinical samples representing both sexes were used.

### Cells

Primary human astrocytes (3H Biomedical - SC1800-5) were cultured in astrocyte medium (3H Biomedical, SC1801) supplemented with 2% FBS, 1% PenStrep and astrocyte growth serum (provided by 3H Biomedical). Multiple batches were used throughout the study, each designated by a different letter. As each batch was derived from a distinct individual, they served as independent biological replicates.

iPSC-derived astrocytes have been generated as previously described (15, 16). Briefly, H1 (WA01) hESCs from WiCell Research Institute (WiCell, WI) were plated on Matrigel (Corning)-coated 6-well plates and infected with 2 µL of each virus (rtTA, Sox9 and Nfib) in mTeSR1 supplemented with 8 µg/mL of protamine sulfate. Twenty-four hours later, medium was changed and fresh mTeSR1 supplemented with 2.5 µg/mL doxycycline was added. Subsequently, selection was conducted for 5 days (2.5 µg/mL Puromycin and 250 µg/mL Hygromycin) days in expansion medium (DMEM F-12, 10% FBS, N2, 1% Glutamax from Thermo Fisher Scientific). At day 7 after induction, cells are plated on Matrigel-coated glass coverslips in 12-well plates and the media is changed to FGF medium (Neurobasal, 2% B27 supplement, 1% NEAA, 1% Glutamax, and 1% FBS, from Thermo Fisher Scientific; 8 ng/mL FGF, 5 ng/mL CNTF, and 10 ng/mL BMP4, from Peprotech). Two days after, FGF medium was replaced, and afterward 50% of the medium was replaced with Maturation medium (1:1 DMEM/F-12 and Neurobasal, 1% N2, 1% sodium pyruvate, and 1% Glutamax, from Thermo Fisher Scientific; 5 mg/mL N-acetyl-cysteine, 500 mg/mL dbcAMP, from Sigma-Aldrich; 5 ng/mL heparin-binding EGF-like growth factor, 10 ng/mL CNTF, 10 ng/mL BMP4, from Peprotech) every 2–3 d, and cells were kept for 14 days before experiment.

The glioma cell lines U3082MG, U3065MG and U3020MG were obtained from Human Glioblastoma Cell Culture resource (hgcc.se) and cultured in HGC medium on laminin-coated plates, as described previously (17). Cells were passaged using Accutase (Thermo Fisher Scientific).

U251MG (MilliporeSigma) and T98G (ATCC) glioma cells were cultured in DMEM supplemented with 10% FBS and 1% PenStrep. The Non-Small Cell Lung Cancer (NSCLC) H2030_BrM3 cells were cultured in RPMI supplemented with 2mM Glutamine, 10% FBS and 1% PenStrep.

Primary murine brain tumor cells were isolated as previously described (18).

All cell lines were tested for mycoplasma contamination every 6 months (MycoplasmaCheck, Eurofins). Cells were maintained at 37°C in a humidified incubator with 21% O2 and 5% CO2, unless otherwise stated.

Irradiation was performed using a Cix1 x-ray irradiator (Xstrahl) at indicated dose. Hypoxic conditions were generated using either a Whitley H35 Hypoxystation (Don Whitley Scientific) or an InvivO2 400 Hypoxia Workstation (Baker Ruskinn) A full list of reagents used for cell treatments is provided in **Table 4**.

### Immunofluorescence and analysis of data

Cells were treated as described in the figure legends and fixed 48 hours after irradiation, temozolomide treatment, or hypoxia induction using 4% paraformaldehyde (PFA) for 20 minutes. After fixation, cells were washed and permeabilized with 0.3% Triton X-100 (Sigma-Aldrich) in PBS. Blocking was performed using 2.5% fish gelatin (Sigma-Aldrich) in PBS containing 0.05% Tween-20 (Sigma-Aldrich), followed by overnight incubation at 4°C with primary antibodies (**Table 2**) prepared in blocking solution. Cells were then incubated with appropriate secondary antibodies (**Table 3**) mixed with phalloidin where applicable at room temperature. Hoechst staining was performed during last PBS wash at 0.5ug/mL. After a final wash with water, coverslips were dried and mounted using ProLong Glass Antifade Mountant (Invitrogen). All the other reagents used for cell treatment or staining are listed in **Table 4**. Images were acquired using an Olympus BX63 microscope equipped with DP80 camera and cellSens Dimension v1.12 software (Olympus Corporation). CellProfiler (www.cellprofiler.org) (19) was used to extract 35 features related to morphology, size and intensity (**Table 1**). Data were analyzed in R, and Principal Component Analysis (PCA) was performed using the FactoMineR and factoextra packages. Intensity and morphology features were represented as Superplots (20).

**Table 1:** variables extracted using CellProfiler and used for PCA.

| Cell Compartment | Variable family | Variable |
| --- | --- | --- |
| Nucleus | Area/Shape | BoundingBoxArea |
| Nucleus | Area/Shape | CentralMoment_0_0 |
| Nucleus | Area/Shape | Eccentricity |
| Nucleus | Area/Shape | FormFactor |
| Nucleus | Area/Shape | HuMoment_0 |
| Nucleus | Area/Shape | InertiaTensorEigenvalues_0 |
| Nucleus | Area/Shape | InertiaTensor_0_0 |
| Nucleus | Area/Shape | SpatialMoment_0_0 |
| Nucleus | Area/Shape | Zernike_0_0 |
| Nucleus | Intensity (Hoechst) | MeanIntensity |
| Nucleus | Intensity (Hoechst) | MassDisplacement |
| Nucleus | Intensity (Hoechst) | StdIntensity |
| Cell | Area/Shape | Area |
| Cell | Area/Shape | CentralMoment_0_0 |
| Cell | Area/Shape | Compactness |
| Cell | Area/Shape | EquivalentDiameter |
| Cell | Area/Shape | InertiaTensorEigenvalues_1 |
| Cell | Area/Shape | MajorAxisLength |
| Cell | Area/Shape | Perimeter |
| Cell | Area/Shape | SpatialMoment_0_0 |
| Cell | Area/Shape | Zernike_2_0 |
| Cell | Intensity (GFAP) | MeanIntensity |
| Cell | Intensity (GFAP) | StdIntensity |
| Cell | Intensity (Phalloidin) | IntegratedIntensity |
| Cell | Intensity (Phalloidin) | MassDisplacement |
| Cell | Intensity (Phalloidin) | MeanIntensityEdge/MeanIntensity |
| Cell | Intensity (Vimentin) | IntegratedIntensity |
| Cell | Intensity (Vimentin) | MeanIntensity |
| Cytoplasm | Texture (Phalloidin) | Correlation_12_00_256 |
| Cytoplasm | Texture (Phalloidin) | InfoMeas1_12_00_256 |
| Cytoplasm | Texture (Phalloidin) | InfoMeas2_12_00_256 |
| Cytoplasm | Texture (Vimentin) | Correlation_12_00_256 |
| Cytoplasm | Texture (Vimentin) | Entropy_Vimentin_12_00_256 |
| Cytoplasm | Texture (Vimentin) | InfoMeas1_Vimentin_12_00_256 |
| Cytoplasm | Texture (Vimentin) | SumVariance_Vimentin_12_00_256 |

**Table 2:**
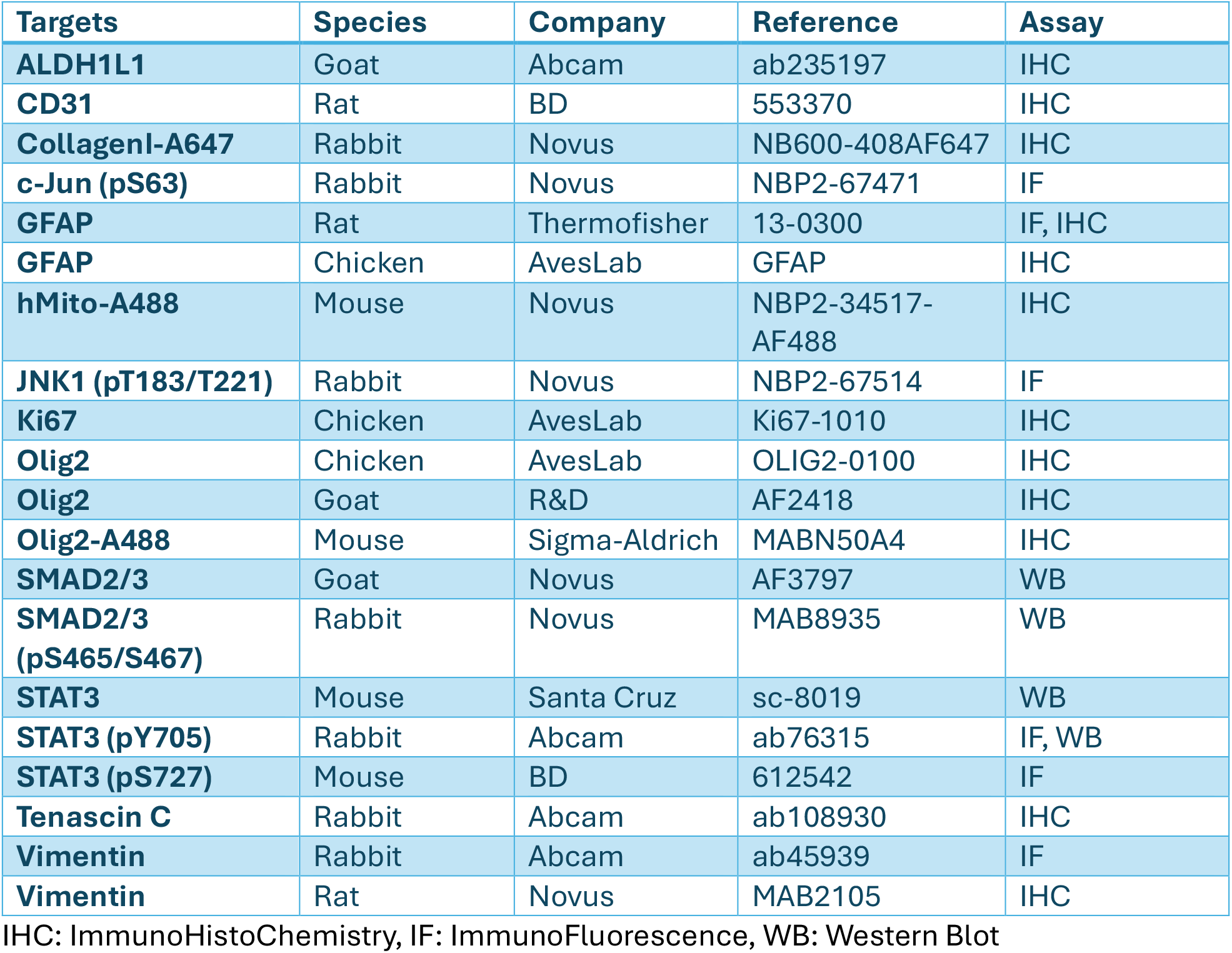
primary antibodies used in the study.

| Targets | Species | Company | Reference | Assay |
| --- | --- | --- | --- | --- |
| **ALDH1L1** | Goat | Abcam | ab235197 | IHC |
| **CD31** | Rat | BD | 553370 | IHC |
| **CollagenI-A647** | Rabbit | Novus | NB600-408AF647 | IHC |
| **c-Jun (pS63)** | Rabbit | Novus | NBP2-67471 | IF |
| **GFAP** | Rat | Thermofisher | 13-0300 | IF, IHC |
| **GFAP** | Chicken | AvesLab | GFAP | IHC |
| **hMito-A488** | Mouse | Novus | NBP2-34517-AF488 | IHC |
| **JNK1 (pT183/T221)** | Rabbit | Novus | NBP2-67514 | IF |
| **Ki67** | Chicken | AvesLab | Ki67-1010 | IHC |
| **Olig2** | Chicken | AvesLab | OLIG2-0100 | IHC |
| **Olig2** | Goat | R&D | AF2418 | IHC |
| **Olig2-A488** | Mouse | Sigma-Aldrich | MABN50A4 | IHC |
| **SMAD2/3** | Goat | Novus | AF3797 | WB |
| **SMAD2/3 (pS465/S467)** | Rabbit | Novus | MAB8935 | WB |
| **STAT3** | Mouse | Santa Cruz | sc-8019 | WB |
| **STAT3 (pY705)** | Rabbit | Abcam | ab76315 | IF, WB |
| **STAT3 (pS727)** | Mouse | BD | 612542 | IF |
| **Tenascin C** | Rabbit | Abcam | ab108930 | IHC |
| **Vimentin** | Rabbit | Abcam | ab45939 | IF |
| **Vimentin** | Rat | Novus | MAB2105 | IHC |

IHC: ImmunoHistoChemistry, IF: ImmunoFluorescence, WB: Western Blot

**Table 3:**
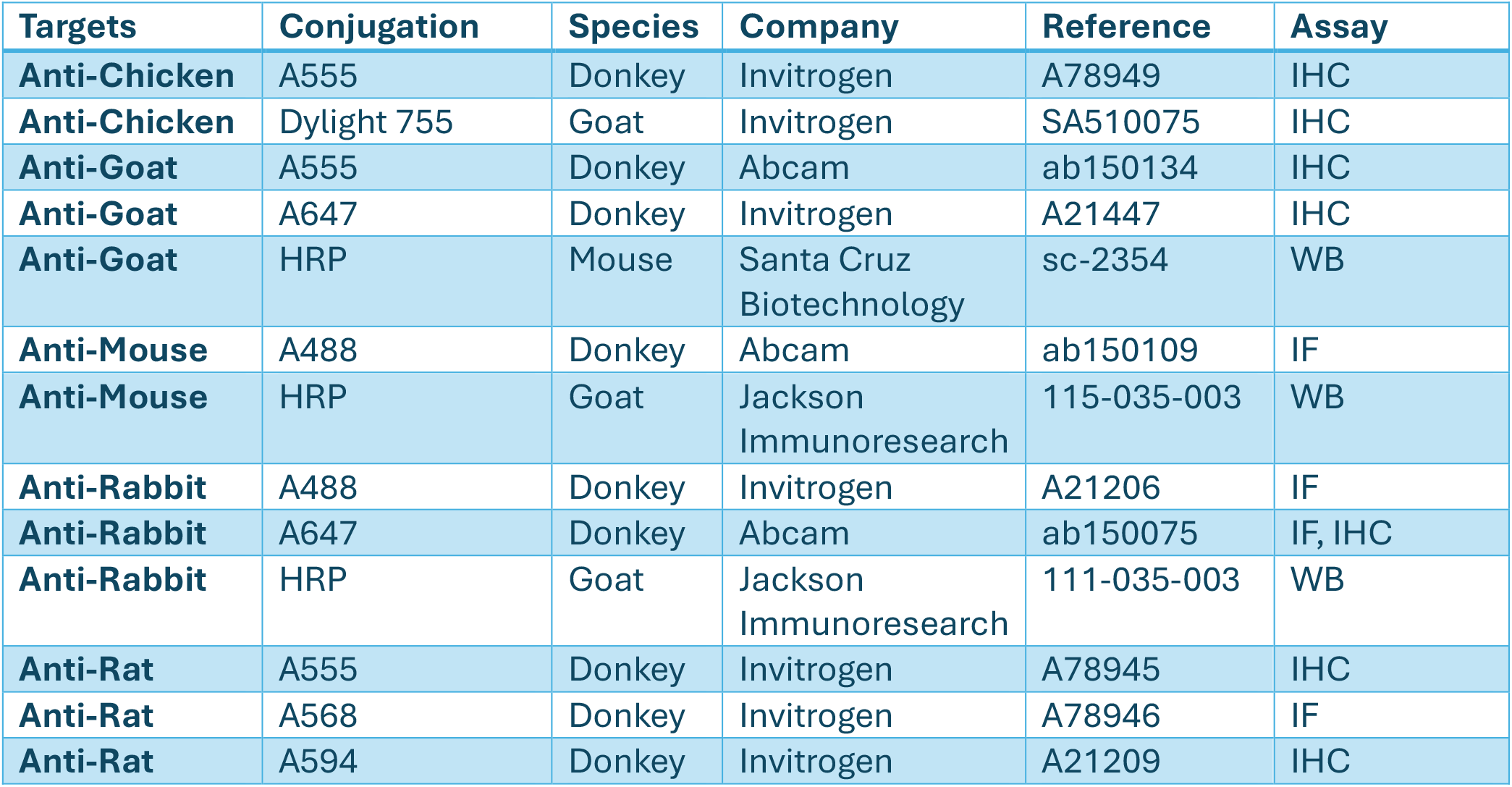
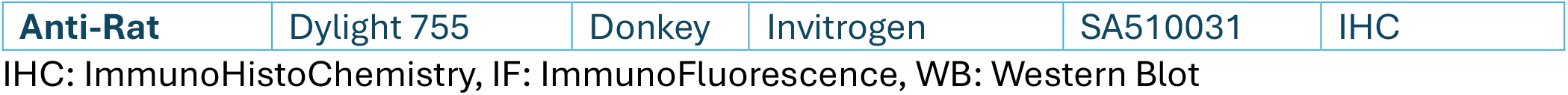
secondary antibodies used in the study.

| Targets | Conjugation | Species | Company | Reference | Assay |
| --- | --- | --- | --- | --- | --- |
| **Anti-Chicken** | A555 | Donkey | Invitrogen | A78949 | IHC |
| **Anti-Chicken** | Dylight 755 | Goat | Invitrogen | SA510075 | IHC |
| **Anti-Goat** | A555 | Donkey | Abcam | ab150134 | IHC |
| **Anti-Goat** | A647 | Donkey | Invitrogen | A21447 | IHC |
| **Anti-Goat** | HRP | Mouse | Santa Cruz Biotechnology | sc-2354 | WB |
| **Anti-Mouse** | A488 | Donkey | Abcam | ab150109 | IF |
| **Anti-Mouse** | HRP | Goat | Jackson ImmunoResearch | 115-035-003 | WB |
| **Anti-Rabbit** | A488 | Donkey | Invitrogen | A21206 | IF |
| **Anti-Rabbit** | A647 | Donkey | Abcam | ab150075 | IF, IHC |
| **Anti-Rabbit** | HRP | Goat | Jackson ImmunoResearch | 111-035-003 | WB |
| **Anti-Rat** | A555 | Donkey | Invitrogen | A78945 | IHC |
| **Anti-Rat** | A568 | Donkey | Invitrogen | A78946 | IF |
| **Anti-Rat** | A594 | Donkey | Invitrogen | A21209 | IHC |
| **Anti-Rat** | Dylight 755 | Donkey | Invitrogen | SA510031 | IHC |

IHC: ImmunoHistoChemistry, IF: ImmunoFluorescence, WB: Western Blot

**Table 4:** reagents used in the study.

| **Reagents** | **Reference** | **Company** | **Assay** |
| --- | --- | --- | --- |
| **DMSO** | D2438 | Sigma-Aldrich | IF, *In vivo* experiments, WB, TOX, CFA |
| **Flunarizine dihydrochloride** | HY-B0358A | MedChemExpress | IF, *In vivo* experiments, WB, TOX, CFA |
| **Hoechst 33342** | B2261 | Sigma-Aldrich | IF, IHC |
| **JSH-23** | 15036 | Cayman Chemical | IF |
| **Neocarzinostatin** | N9162 | Sigma-Aldrich | IF |
| **Phalloidin iFluor 488** | ab176753 | Abcam | IF |
| **Phalloidin iFluor 647** | 20555 | Cayman Chemical | IF |
| **Silibinin** | HY-13748 | MedChemExpress | IF |
| **SP600125** | 10010466 | Cayman Chemical | IF |
| **SR-3306** | HY-12829 | MedChemExpress | IF |
| **Temozolomide** | T2577 | Sigma-Aldrich | IF |
| **TGF-β1 (human recombinant)** | 100-21 | Peprtech | WB |
| **TUNEL** | 11767305001 (Enzyme) + 11767291910 (Label) | Roche | IHC |

IHC: ImmunoHistoChemistry, IF: ImmunoFluorescence, WB: Western Blot, TOX: Toxicity assay, CFA: Colony Formation Assay

### Phospho-antibody array

The Phospho Explorer antibody array (PEX100, Full Moon Biosystems) was performed according to the manufacturer’s protocol using primary human astrocytes, either irradiated or non-irradiated. Slides were scanned using an INNOPSYS InnoScan Microarray scanner and signal intensity data for each spot on the array were extracted from the images. Mean signal intensity values were calculated from duplicate spots. Phospho-protein signals were normalized to their corresponding total protein signals, and fold changes were determined by comparing irradiated samples to non-irradiated controls.

### Drug screening

For the primary drug screens, primary human astrocytes from two different batches were seeded in 384-well plates and cultured as described above. On the second day, cells were treated with one of the following libraries: 176 compounds from Enzo SCREEN-WELL Protease, Kinase and Epigenetics (PKE) Inhibitor libraries, 330 compounds from the TargetMol Anti-Cancer drugs library (#L2110, v2018) or 1,280 compounds from the FDA-approved Prestwick Chemical library (prestwickchemical.com, v2016). Compounds were tested at concentrations of 1µM and 3µM (PKE library & TargetMol libraries) or 1µM and 10µM (Prestwick Chemical library). Two hours after compound addition, cells were irradiated with 10 Gy. Forty-eight hours post-irradiation, cells were fixed with 4% PFA and immunolabelled as described in the “Immunofluorescence and Analysis of Data” section to detect GFAP, vimentin, Hoechst, and phalloidin signals. Images were acquired using high-throughput microscope (IN Cell Analyzer 2200 for the PKE and Prestwick libraries; ImageXpress Confocal HT for the TargetMol library). CellProfiler v3.0-4.0 (19) was used for nuclear and cytoplasmic segmentation. Between 378 and 463 variables were extracted and used in subsequent analysis (the list of variables used for each drug screen is available in **Supp. Table 1**). Briefly, the pipeline used Hoechst signal to identify nuclei and GFAP, vimentin and phalloidin signals to define cytoplasm regions. Morphological features (e.g., area, perimeter, shape descriptors), intensity features (for all four markers) and texture features (for vimentin and phalloidin) were quantified for nuclei, cytoplasm or whole cells. Quality control was performed using CellProfiler Analyst v3.0 (21) following established recommendations (22). Furthermore, wells with a cell count *z*-score greater than 1.5 were excluded from the analysis to eliminate compound–concentration pairs that were toxic to astrocytes. Quantitative multivariate datasets were exported and then analyzed using HC StratoMineR (Core Life Analytics) (23). Redundant variables were excluded from analysis, and data were normalized across plate and scaled. Principal Component Analysis (PCA) was performed using generalized weighted least squares, and a distance score was computed based on similarity to the non-irradiated control using Pearson approach.

For the confirmation screens, the same method was applied but using four batches of primary astrocytes, four concentrations (0.25 µM, 1 µM, 3 µM and 10 µM) and duplicate wells. Compounds included were those identified as hits in both astrocyte batches during the primary screens, along with 15 manually selected compounds. One astrocyte batch (AstroJ) was excluded from the mean distance score calculation due to its divergent response compared to the other three batches.

### Western blot

Whole cell lysates were prepared in RIPA buffer supplemented with a protease inhibitor cocktail (cOmplete - Roche) and phosphatase inhibitor (PhosSTOP – Roche). Protein concentrations were determined using the Bradford assay. Equal amounts of protein were mixed with NuPAGE LDS Sample Buffer (Invitrogen) and dithiothreitol (DTT; Sigma-Aldrich), then boiled at 70°C for 10 minutes. Proteins were separated on 4–20% Mini-PROTEAN TGX Stain-Free Protein Gels (Bio-Rad Laboratories) and transferred to PVDF membranes using the Trans-Blot Turbo Transfer System (Bio-Rad Laboratories). Membranes were blocked in 5% non-fat dry milk in TBS-T (TBS with Tween 20) and incubated overnight at 4°C with primary antibodies (**Table 2**) diluted in blocking solution or in 5% BSA (Sigma-Aldrich) in TBS-T for phospho-protein detection. After washing, membranes were incubated for 1 hour with secondary antibodies (Jackson ImmunoResearch) (**Table 3**) at room temperature. Signal detection was performed using an Amersham Imager 600 (GE HealthCare). Band intensities were quantified using Fiji and phospho-protein levels were normalized to the corresponding total protein levels.

### Viability assay

Cells were seeded in 96-well plates and treated with or without Flunarizine dihydrochloride. Irradiation was performed 2 hours after drug treatment where applicable. After 3 days, cell viability was assessed by adding WST-1 reagent (ab155902, Abcam) to the culture medium. Absorbance was measured at 450 nm in a Synergy 2 plate reader (BioTek). The mean absorbance of triplicates was calculated, and fold changes in viability were determined by comparing each condition to the non-irradiated, DMSO-treated control.

### Colony formation assay

Cells were seeded at clonal density in 6-well or 12-well plates. On day 2, treatments and irradiation were applied as described in the figure legends. Cells were then allowed to form colonies for 7 to 14 days depending on the cell type. Colonies were fixed in 4% PFA, stained with 0.01% Crystal Violet and manually counted. The mean colony number from triplicates was calculated.

### RNA-seq

RNA extraction was performed from primary human astrocytes 24 hours after they received treatment and irradiation using RNeasy mini kit (Qiagen) together with the Qiashredder Kit (Qiagen) according to manufacturer’s protocols. The sequencing was performed at the Center for Translational Genomics (Lund University). The mRNA library was prepared using Illumina® Stranded mRNA Prep Ligation kit (20091659, Illumina). The sequence was performed using NovaSeq 6000 System (Illumina) and NovaSeq 6000 S4 Reagent Kit v1.5 (20028312, Illumina). To process and analyze the data, paired-end RNA-seq data from irradiated and Flunarizine-treated samples were processed using the nf-core/rnaseq pipeline (v3.14.0) (24), utilizing Salmon for transcript quantification (25). Gene-level expression was aggregated with tximport (26), and the resulting count matrix was loaded into R as a DESeq2 object (27). Gene annotations, including external gene symbols and biotypes, were retrieved from biomaRt (28) and merged into the DESeq2 object. Protein coding genes with a minimum total count greater than 1 across samples were kept. For global differential expression analysis, expression data were modeled using a generalized linear model in DESeq2 with the formula ‘~ Rep + Irradiation + Treatment’. Variance-stabilizing transformation (VST) was applied to normalized counts for PCA, visualizing sample clustering according to treatment and irradiation conditions (**Supp. Fig. 7C**). Calcium signaling gene signatures (“GOBP_POSITIVE_REGULATION_OF_CALCIUM_MEDIATED_SIGNALING” and “GOBP_CALMODULIN_DEPENDENT_KINASE_SIGNALING_PATHWAY”) were obtained from MSigDB via msigdbr (29). Heatmaps representing gene signature expression were generated from median-centered normalized counts (**Supp. Fig. 7A-B**). Differential expression and pathway analyses were performed separately for Flunarizine effect vs DMSO in irradiated cells (IR + Flunarizine vs IR + DMSO, n=8) and for the irradiation effect in untreated cells (IR + DMSO vs nonIR + DMSO, n=8). Differential expression statistics from DESeq2 were used for pathway activity inference with decoupleR (30) and the PROGENy network (31). Pathway activities were estimated using a multivariate linear model and visualized as bar plots (**Fig. 3G**). For IR + Flunarizine vs IR + DMSO, heatmap of the top and bottom 32 significantly differentially expressed genes (by DESeq2 statistics) were created using regularized log-transformed (rlog) expression values to visualize differential gene expression between IR + Flunarizine and IR + DMSO conditions (**Fig. 3E**).

All raw data is available at Array Express E-MTAB-15215.

### Cytokine array

The Human Cytokine Antibody Array (ab133997, Abcam) was performed according to manufacturer’s instructions using 200μg of lysates from primary human astrocytes, either irradiated or non-irradiated. In accordance with the manufacturer’s protocol, data were normalized to the array’s positive control, the non-irradiated control, and total protein concentration. Results are presented as fold changes comparing irradiated samples to non-irradiated controls.

### In vivo models

All animal procedures were conducted in accordance with the European Union directive on animal welfare, and procedures were approved by the regional ethics committee Malmö - Lunds Djurförsöksetiska Nämnd (M-16123/19).

Gliomas were induced in *Nestin-tv-a* mice by intracranial injection RCAS-PDGFB- and RCAS-shp53– transfected chicken fibroblast DF-1 cells (ATCC CRL-12203, ATCC) into the neonatal brain, as previously described (32).

For the survival cohort, mice exhibiting glioma symptoms or signs of hydrocephaly prior to or within day 28 after intracranial injection were euthanized and excluded from the study. Whole brain irradiation was administered on day 28 after intracranial injection. Mice were sedated with isoflurane, and a single dose of 10 Gy was delivered using a 10 mm field on a 220 kV preclinical irradiator (XenX, XStrahl Inc.). Mice received intraperitoneal injections of either vehicle (6% DMSO, 40% PEG300, 5% Tween80, 49% PBS) or Flunarizine dihydrochloride (30mg/kg in vehicle) prepared in vehicle. Mice received treatments for 3 days before and 4 days after irradiation. Subsequently, treatments were administered on a 5-days-on/2-days-off schedule for a maximum of 30 total injections. Mice were monitored daily and euthanized upon the development of glioma symptoms or upon reaching the endpoint of 100 days after intracranial injections. The Kaplan–Meier survival curve was generated using the R package survminer (33). The compounds used to treat the mice are listed in **Table 4**.

For the post-irradiation cohort, a similar protocol was followed, except that mice were treated only for 3 days before and 3 days after irradiation. All mice were euthanized on day 31 after intracranial injections. Animals exhibiting glioma symptoms before day 31 were euthanized and excluded from analysis.

For the brain metastasis cohort, athymic *Foxn1*^nu/nu^ pups (Charles River) were injected intracranially with 5,000 cells from the brain-metastatic derivative of the human lung adenocarcinoma cell line H2030 (H2030_BrM3; provided by Joan Massagué, MSKCC) (34). Mice received irradiation and treatment as described for the survival cohort and were euthanized upon the development of brain tumor symptoms or after losing more than 10% of body weight within one week.

### Tissue staining and analysis

Whole mouse brains were embedded in Optimal Cutting Temperature (OCT) compound (Thermo Fisher Scientific), frozen and cryosectioned. Sections were air-dried for 10 min and fixed in ice-cold acetone. Permeabilization was performed using 0.3% Triton X-100 in PBS, following by blocking in 2.5% fish gelatin with 0.05% Tween20 in PBS. Sections were then incubated overnight at 4°C with primary antibodies (**Table 2**) diluted in blocking solution. After washing, sections were incubated with appropriate secondary antibodies (**Table 3**) at room temperature. Hoechst staining was performed during the last PBS wash at 0.5ug/mL. Sections were mounted using Vectashield Vibrance Antifade Mounting Medium (Vector Laboratories). All the additional reagents used to stain the sections are listed in **Table 4**. Stained sections were imaged using a PhenoImager HT instrument (Akoya Bioscience). Spectral unmixing of images was performed using the inform software package (Akoya Biosciences). Analysis of the pictures was conducted using QuPath (35).

### Patient cohort analysis

Data from Chinese Glioma Genome Atlas (CGGA) (36) were obtained using the Gliovis data portal (https://gliovis.bioinfo.cnio.es) (37) and figures were prepared using R, R studio as well as the R package *survminer* to generate the Kaplan-Meier survival curves.

### Statistics

Unless otherwise stated, all values represent mean ± SD for at least three biological replicates. All statistical analyses were performed in R using RStudio and ggpubr package. Comparisons between two groups were conducted using unpaired Student’s *t*-test unless otherwise specified. For multiple comparisons, one-way ANOVA followed by post-hoc *t*-tests was used when the ANOVA *P*-value was <0.05. For survival curves, *P* values were obtained with Gehan-Breslow (generalized Wilcoxon) tests, followed by log-rank tests to compare groups unless otherwise stated. Three levels of significance were used. A *P*-value of less than 0.05 were considered significant.

### Illustration and Writing Tools

Schematic Illustrations describing the drug screening workflow (**Fig. 2A**), *in vivo* studies design (**Fig. 4A, 4E, 4H, 4L**), the summary of findings (**Fig. 6**) and the graphical abstract were created using BioRender.com. All figures have been edited using Affinity Designer v2.6.2. Language editing and grammar corrections were assisted by ChatGPT-4 (OpenAI).

## Results

### A novel image-based approach to monitor astrocyte reactivity

Our first objective was to develop a method to monitor astrocyte reactivity that relied not on a single marker, as discouraged by a recent consensus statement (5), but instead on a combination of morphological features and reactive astrocyte marker expression. Following IR, we observed significant alterations in both cell and nuclear morphology, as well as increased expression of established reactive astrocyte markers such as GFAP and vimentin, in both primary human astrocytes and iPSC-derived astrocytes (**Fig. 1A**). Some of these changes are well-recognized hallmarks of astrocyte reactivity, including enlarged cell body (**Fig. 1B**), increased GFAP expression (**Fig. 1C**) and vimentin expression (**Fig. 1D**) (5). Additional morphological features were also affected by IR, including nuclear and cell shape descriptors and phalloidin signal intensity (**Supp. Fig. 1 A-E**).

**Figure 1:**
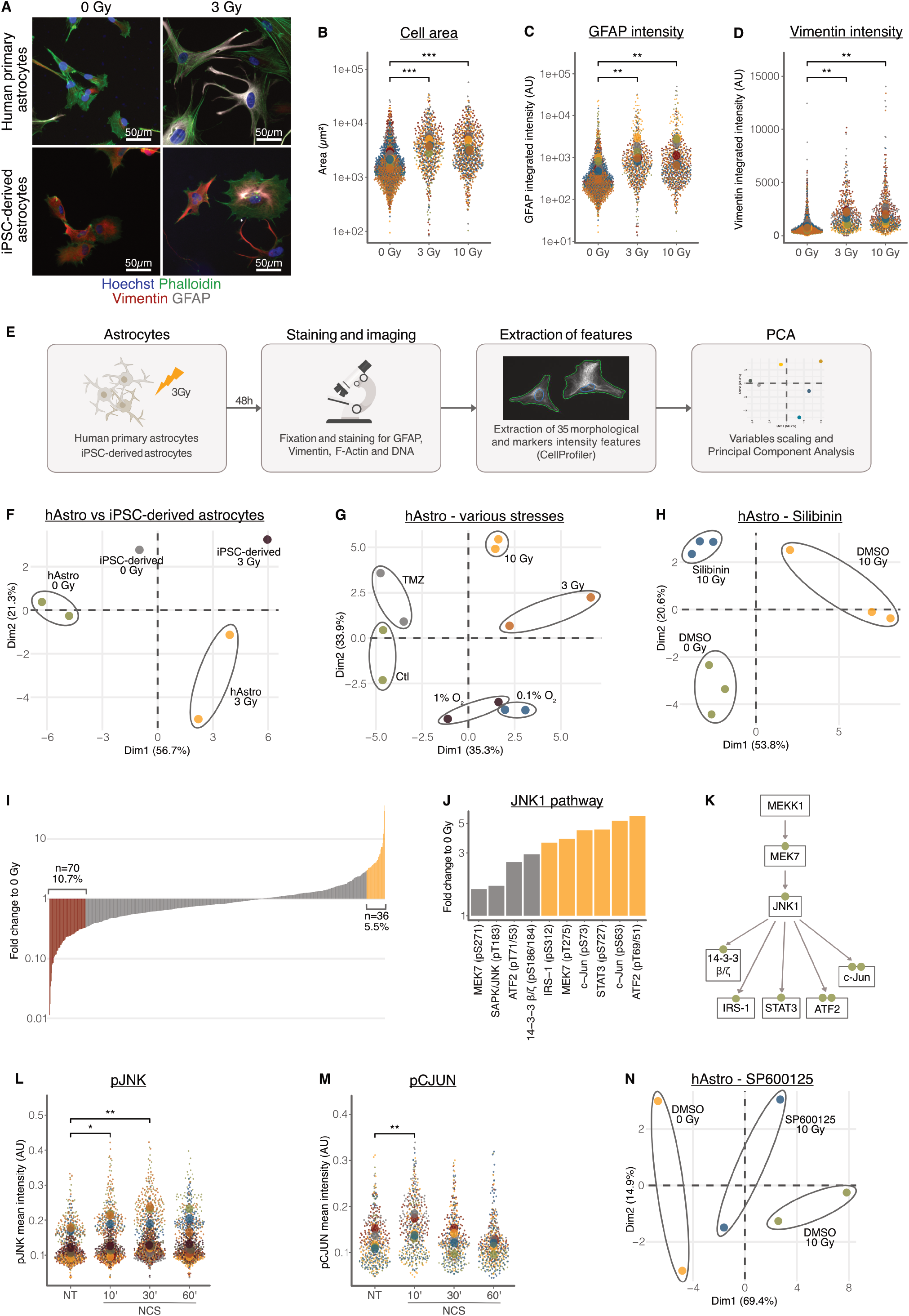
An image-based approach to monitor astrocyte reactivity allows the identification of JNK pathway as required for the induction of reactivity by IR. **(A)** Representative images of immunofluorescent detection of DNA (Hoechst-33342), F-Actin (Phalloidin), vimentin and GFAP in primary human astrocytes and iPSC-derived astrocytes with or without 3 Gy irradiation. **(B-D)** Quantification of cell area **(B)**, GFAP integrated intensity **(C)** and vimentin integrated intensity **(D)** in primary human astrocytes irradiated with 0, 3 or 10 Gy and immunofluorescent staining as described for **(A)** (*n*=3). **(E)** Flowchart describing the phenotypic analysis pipeline used on astrocytes. **(F-H)** Principal component analysis (PCA) of the morphological and intensity features extracted as described in **(E)** comparing: response to IR of primary human astrocytes (hAstro) vs. iPSC-derived astrocytes (iPSC-derived) **(F)**; response of primary human astrocytes to various stresses vs. control (Ctl) (0 Gy, 21% O_2_) **(G)**; and response of primary human astrocytes to IR and STAT3i Silibinin (10µM) **(H). (I-J)** Phospho-protein array performed on primary human astrocytes 1 hour post irradiation with 10 Gy IR detecting phosphorylation levels on 1,318 phosphosites. Results are shown as fold change relative to non-irradiated controls. Bars are colored red if fold change <3, yellow if >3, and gray otherwise. Figures show all the phosphosites **(I)** or only the ones involved in the canonical JNK pathway **(J)** described in **(K). (L-M)** Quantification of pT183/Y185-JNK1/2/3 (*n*=9) **(L)** or pS63-CJUN (*n*=8) **(M)** nuclear mean intensities following immunofluorescent detection in primary human astrocytes treated or not with the radiomimetic compound Neocarzinostatin (NCS) (100ng/mL) for 10, 30 or 60 minutes. **(N)** PCA of morphological and intensity features extracted as described in **(E)** comparing response to IR and JNKi SP600125 (10µM) in primary human astrocytes. **(B-D, L-M)** Results are represented as SuperPlots, with small dots representing individual measurements and large dots indicating means per biological replicate. Each color indicates a distinct biological replicate. Fluorescence intensity is shown in arbitrary units (AU). Experiments analyzed by PCA were repeated at least three times; one representative result is shown. *P*-values were calculated using paired *t-*tests. \**P* < 0.05, \*\**P* < 0.01, \*\*\**P* < 0.001.

To comprehensively capture these changes, we selected 35 quantifiable features that were the most altered in astrocytes after IR (**Table 1**), using CellProfiler (19). We then applied Principal Component Analysis (PCA) to integrate and analyze these variables, providing a multidimensional representation of astrocyte reactivity (**Fig. 1E**). Using this approach, we successfully detected changes associated with reactivity in both primary human astrocytes and iPSC-derived astrocytes following treatment with 3 Gy IR. Interestingly, both cell types exhibited similar variance changes along the first principal component (Dimension 1), suggesting a conserved reactivity response to IR across different astrocyte models (**Fig. 1F**). Next, we exposed astrocytes to different stress conditions relevant to the GBM microenvironment, namely temozolomide, representing standard chemotherapeutic treatment, and hypoxia, which is characteristic of the GBM tumor niche (**Fig. 1G**). In primary human astrocytes, temozolomide exposure induced only a mild effect, whereas both IR (3 and 10 Gy) and hypoxia caused pronounced alterations in morphology and marker expression, with more marked effects observed at 0.1% O_2_ compared to 1% O_2_. These findings are consistent with our previous work showing that hypoxia induces astrocyte reactivity with a more robust phenotype at 0.1% O_2_ than at 1% O_2_ (38). However, in iPSC-derived astrocytes, we only observed a modest effect of hypoxia at 1% O_2_ while 0.1% O_2_ induced strong changes (**Supp. Fig. 2A**). Overall, iPSC-derived astrocytes responded strongly at 0.1% O_2_ and 3 Gy IR but showed only mild changes after temozolomide treatment and 10 Gy IR (**Supp. Fig. 2B**).

To validate our approach, we tested the effect of the STAT3 inhibitor Silibinin, which has been shown to counteract the tumor-promoting effects of astrocytes in the brain TME (12). We first confirmed STAT3 activation by detecting increased phosphorylation of Y705-STAT3 following IR in primary human astrocytes (**Supp. Fig. 2C**). Co-treatment with Silibinin partially reversed the IR-induced phenotype (**Fig. 1H**). Similarly, inhibition of the NF-κB pathway - another key mediator of astrocyte reactivity - also resulted in a partial rescue of the IR-induced phenotype (**Supp. Fig. 2D**). Together, these results establish a robust, multidimensional image-based approach to quantify astrocyte reactivity, capable of capturing diverse reactive phenotypes and their modulation by environmental stressors or pharmacological interventions.

### The JNK pathway is required for the induction of IR-induced astrocyte reactivity

To identify signaling pathways essential for IR-induced astrocyte reactivity, we first performed a phospho-protein array including 1,318 phosphosites that compared phosphorylation levels between astrocytes treated with 10 Gy IR and non-irradiated controls (**Fig. 1I, Supp. Fig. 3A-B, Supp. Table 2**). This analysis revealed increased phosphorylation within the canonical JNK1 pathway (**Fig. 1J–K**), in contrast to other JNK substrates such as those from the Bcl-2 family (**Supp. Fig. 3C**). To confirm JNK pathway activation in response to IR, we treated primary human astrocytes with Neocarzinostatin, a radiomimetic agent that mimics the IR-induced DNA-damage (39). Shortly after Neocarzinostatin treatment, we observed increased phosphorylation of pT183/Y185-JNK1/2/3 (**Fig. 1L**) and pS63-CJUN (**Fig. 1M**), indicating rapid activation of JNK pathway. Additionally, pS727-STAT3, a described JNK1 target site, also displayed increased phosphorylation following IR (**Supp. Fig. 3D)**. To assess whether JNK signaling is functionally required for astrocyte reactivity, we applied our phenotypic approach (**Fig. 1E**) and found that SP600125, a JNK inhibitor, partially reversed the IR-induced phenotype (**Fig. 1N**). A similar rescue effect was observed with SR3306, a highly selective JNK inhibitor developed for therapeutic use (**Supp. Fig. 3E**) (40). Together, these results identify the JNK pathway as a key mediator of IR-induced astrocyte reactivity and validate the potential of our image-based approach as a novel tool to investigate astrocyte reactivity.

### Identification of new compounds inhibiting IR-induced astrocyte reactivity

To identify novel inhibitors of IR-induced astrocyte reactivity, including candidates for drug repurposing, we applied our image-based approach at a larger scale. We screened three different compound libraries: 1) a small library of 176 kinase, protease and epigenetic inhibitors (PKE library); 2) a library of 330 anti-cancer drugs tested in humans (Anti-Cancer library); and 3) a larger library composed of 1,280 FDA-approved compounds (FDA Approved library) (**Fig. 2A**). To minimize batch-specific effects, two independent batches of primary human astrocytes were used (AstroD and AstroF). Cells were treated, fixed, and stained as previously described. We extracted around 400 morphology and intensity features from the images using CellProfiler (**Supp. Table 1**) describing cell and nuclear area, shape, signal intensities and cytoplasmic texture for phalloidin and vimentin signal (**Supp. Fig. 4A**). Image quality was assessed using CellProfiler Analyst, following established guidelines to exclude images with poor focus, saturation artifacts, or segmentation errors (22). Using HC StratoMineR, a web-based tool for high-content data analysis (23), we scaled and normalized the data for each compound library and astrocyte batch, and performed PCA, revealing clear separation between irradiated and non-irradiated controls (**Supp. Fig. 4B-D**). To quantify this separation, a distance score to non-irradiated controls was computed, providing quantitative support for the observed divergence between conditions (**Fig. 2B, Supp. Table 3-5**). Compounds yielding a distance score below 0.7 were considered hits. The proportion of hits detected in one or both astrocyte batches was visualized using pie charts (**Fig. 2C**). Interestingly, hits from the FDA-approved library spanned a wide range of therapeutic classes, with no enrichment for specific classes observed.

**Figure 2:**
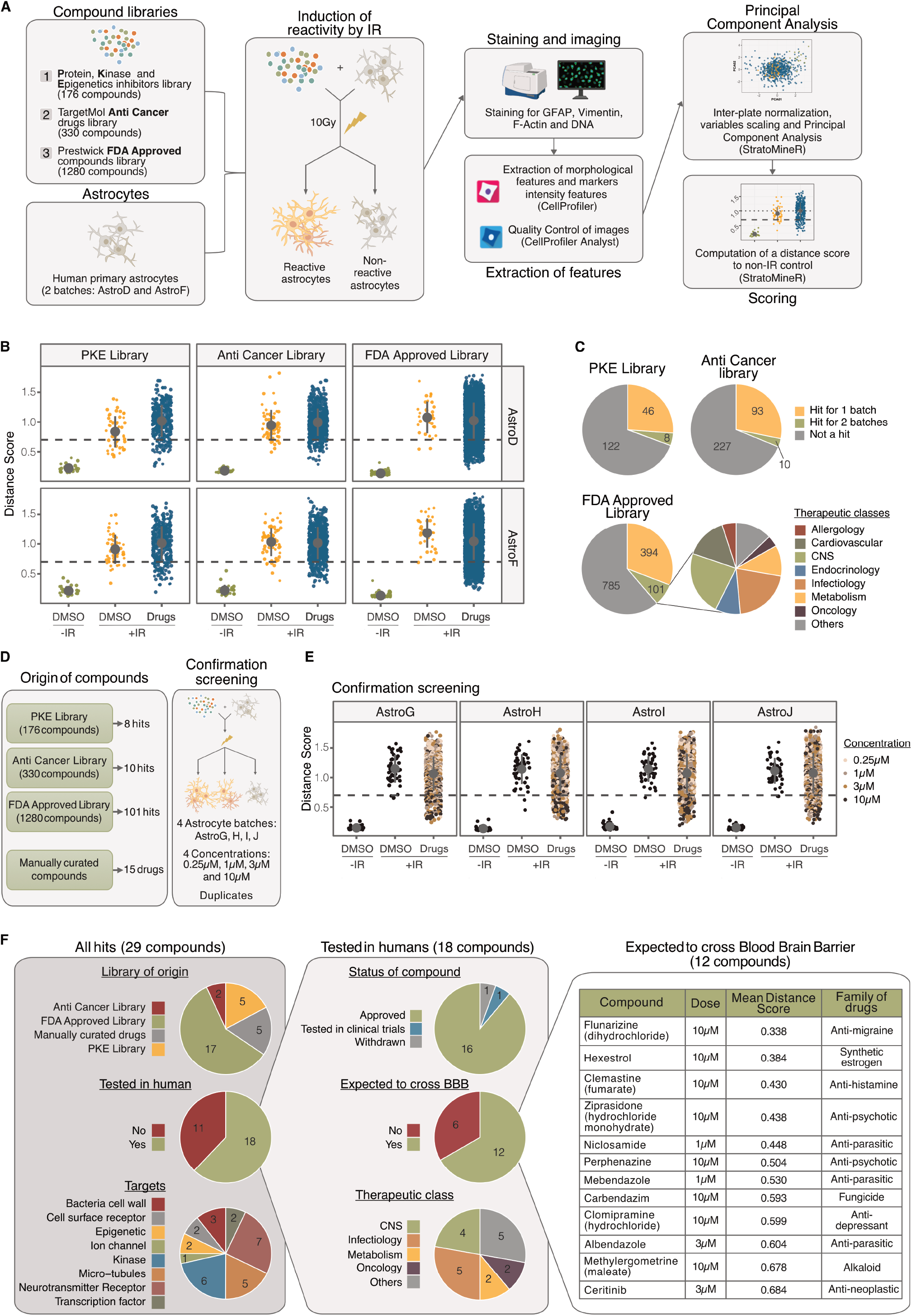
A phenotypic drug screen identifies novel compounds that inhibit IR-induced astrocyte reactivity. **(A)** Flowchart describing the workflow used to identify compounds that inhibit IR-induced astrocyte reactivity in primary human astrocytes. **(B)** Jitter plot showing the results of the primary screens, represented as distance scores to the non-IR control for two primary human astrocyte batches and three compound libraries. The dashed line indicates the hit threshold. **(C)** Pie charts showing the frequency of hits across the three libraries and the distribution of therapeutic classes among hits from the FDA-approved Prestwick Chemicals library. **(D)** Schematic illustrating the origin of compounds and the experimental strategy used for the confirmation screening. **(E)** Jitter plot of confirmation screen results represented as distance score to the non-IR control across four primary human astrocyte batches. The dashed line represents the hit threshold. **(F)** Summary of confirmed hits categorized by compound origin, clinical testing, blood–brain barrier (BBB) permeability, therapeutic class, and molecular targets. The final 12 prioritized compounds are listed.

To validate the initial hits, we performed a confirmation screen that included all compounds scoring as hits in both batches from the primary screens, supplemented with 15 manually selected compounds. These additional candidates were chosen based on prior literature reports or because they target kinases implicated in the phosphorylation of top hits from the phospho-antibody array (**Supp. Fig. 3A**). In total, 134 compounds were tested across four astrocyte batches at four concentrations and in duplicates - substantially increasing data robustness (**Fig. 2D**). PCAs and distance scores again showed good separation for all astrocyte batches between controls (**Supp. Fig. 5A, Fig. 2E, Supp. Table 6**). Mean distance scores were calculated across three batches and a threshold of 0.7 was applied to define final hits (**Supp. Fig. 5B**).

The final hit list included 29 compounds, originating from all three libraries and targeting diverse cell components (**Fig. 2F**.). To prioritize candidates suitable for drug repurposing in GBM, we focused on 18 compounds that had already been tested in humans. Of these, 16 are currently approved drugs across various therapeutic classes. We further refined the list to 12 compounds predicted to cross the blood-brain barrier, a critical feature for Central Nervous System (CNS)-targeted therapies. These included drugs used for migraine, psychosis, and depression, as well as several anti-parasitic agents - accounting for 25% of the final hits. Visual inspection of images confirmed that these compounds inhibited IR-induced morphological changes in astrocytes (**Supp. Fig. 6**). Among the 12 hits, Flunarizine was selected for further investigation. It consistently ranked among the lowest distance scores in both the initial and confirmation screens, suggesting strong efficacy in suppressing astrocyte reactivity. Additionally, Flunarizine is an off-patent drug that has been approved for decades, primarily for the treatment of migraines, and has a well-established safety profile (41, 42), making it a viable candidate for drug repurposing in GBM. Together, these results demonstrate the effectiveness of our high-content, image-based screening approach in identifying clinically relevant compounds that inhibit IR-induced astrocyte reactivity and hold promise for drug repurposing in GBM.

### Flunarizine inhibits *in vitro* IR-induced astrocyte reactivity without toxic eeect

To validate the drug screen results, we repeated immunofluorescence and PCA as described in Fig. 1E. Both the images (**Fig. 3A**) and the PCA (**Fig. 3B**) confirmed that Flunarizine partially reversed IR-induced morphological changes in astrocytes. Phosphorylation of Y705-STAT3 has been described previously as a marker of astrocyte reactivity inducing the tumor-promoting functions of astrocytes (12). Western blot analyses revealed an inhibition of STAT3 phosphorylation upon Flunarizine treatment 15 minutes after IR in comparison with DMSO-treated controls (**Fig. 3C**). To assess potential cytotoxicity, we performed a WST-1 assay in primary human astrocytes and found no toxic effect of Flunarizine, either alone or in combination with IR (**Fig. 3D**).

**Figure 3:**
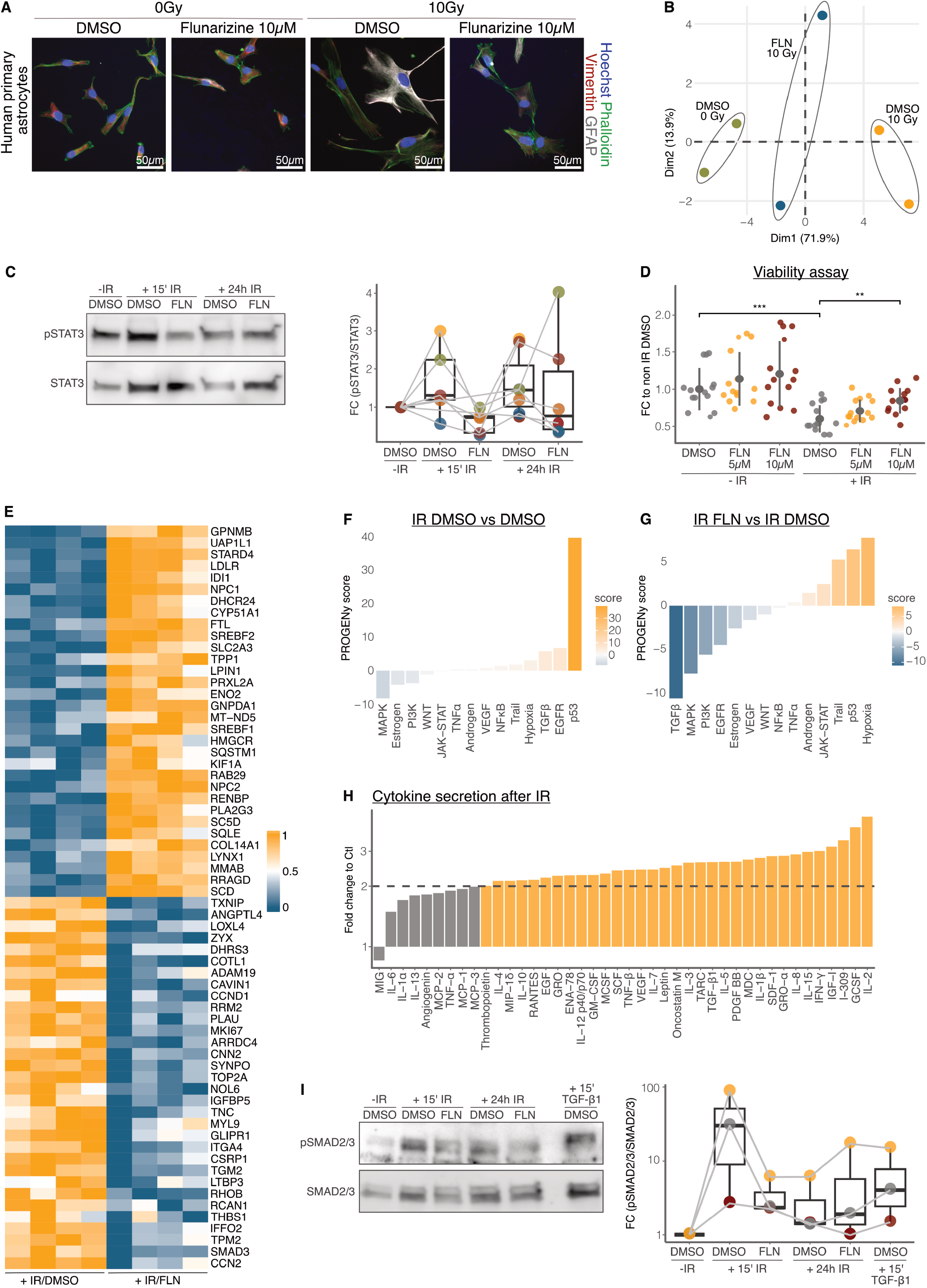
Flunarizine inhibits astrocyte reactivity through suppression of TGF-β pathway. **(A)** Representative images showing immunofluorescent detection of DNA (Hoechst-33342), F-Actin (Phalloidin), vimentin and GFAP in primary human astrocytes treated or not with irradiation and Flunarizine. **(B)** PCA of morphological and intensity features (as described in Figure 1E) comparing combined responses to IR and Flunarizine (FLN) of primary human astrocytes. The experiment was repeated three times; one representative result is shown. **(C)** Immunoblot analysis of pY705-STAT3 and total STAT3 in primary human astrocytes treated with 10 Gy and Flunarizine. Right panel shows the quantification of pSTAT3 signal/STAT3 signal is shown as fold change (FC) relative to the non-IR DMSO control. Each color represents a biological replicate. **(D)** WST-1 assay using primary human astrocytes treated with 10 Gy and Flunarizine. Results are expressed as fold change relative to the mean of non-IR DMSO controls within the same experiment. Data show mean ± SD (*n*=14). *P*-value was determined using one-way ANOVA followed by post-hoc unpaired *t*-tests. \*\**P* < 0.01, \*\*\**P* < 0.001. **(E-G)** RNA sequencing of primary human astrocytes treated with 10 Gy and DMSO or Flunarizine. **(E)** Heatmap showing expression of the top 32 up- and downregulated genes between IR/DMSO and IR/Flunarizine samples (*n*=4). Expression values are rlog-transformed and min– max scaled per gene. **(F-G)** Pathway activity scores inferred using PROGENy and decoupleR for the IR/DMSO vs. nonIR/DMSO data (*n*=4) and IR/Flunarizine vs. IR/DMSO data (*n*=4), respectively. Pathways are ranked by scores derived from the DESeq2 test statistics. **(H)** Cytokine array analysis of primary human astrocytes treated with 10 Gy. The results are shown as fold change relative to non-IR control. The dashed line represents a 2-fold increase. **(I)** Immunoblot analysis of pS465/S467-SMAD2/3 and total SMAD2/3 in primary human astrocytes treated with 10 Gy or TGF-β1 (10ng/mL) and Flunarizine or DMSO treatment. Right panel shows the quantification of (pSMAD2/3 signal)/(SMAD2/3 signal) shown as fold change relative to non-IR DMSO controls (*n*=3). For all Flunarizine treatments in these experiments, Flunarizine dihydrochloride was used at 10 µM unless otherwise specified.

### Flunarizine’s eeect on astrocytes is mediated through inhibition of TGF-β pathway

To elucidate the mechanism-of-action by which Flunarizine inhibits astrocyte reactivity, we performed bulk RNA-sequencing on primary human astrocytes treated with either DMSO or Flunarizine followed by exposure to 0 Gy or 10 Gy IR (**Supp Fig 7C**). Interestingly, by comparing astrocytes that received DMSO or Flunarizine in combination with 10 Gy, among the most downregulated genes, there were several genes coding for extracellular matrix (ECM) proteins (*TGM2, TNC*) or ECM-interacting proteins (*LOXL4, ZYX, ADAM19, LTBP3, ITGA4, THBS1, CCN2*) (**Fig. 3E**). Surprisingly, despite Flunarizine being known as both a calcium channel and calmodulin inhibitor, Flunarizine treatment-alone or in combination with IR-had no detectable effect on the gene signatures associated with “positive regulation of calcium-mediated signaling” or “calmodulin-dependent kinase signaling” (**Supp Fig. 7A-B**). Pathway activity inference using decoupleR (30) and PROGENy (31) revealed that IR and DMSO increased p53 signature activity in comparison to the DMSO control (**Fig. 3F**), as expected following DNA damage, and consistent with the reduced astrocyte viability observed after IR (**Fig. 3D**). Additionally, IR induced significant EGFR and TGF-β pathway activities, both known regulators of astrocyte reactivity (43–45). Notably, co-treatment with Flunarizine significantly suppressed the expression of these signatures (**Fig. 3G**). Moreover, cytokine array analysis showed increased secretion of both TGF-β1 and EGF among the many cytokines secreted following exposure to 10 Gy IR, among other induced cytokines (**Fig. 3H**). Finally, western blot analysis demonstrated that IR induced phosphorylation of SMAD2 and SMAD3, key downstream effectors of the TGF-β pathway, which was effectively inhibited by Flunarizine 15 minutes after IR exposure (**Fig. 3I**). Together, these findings demonstrate that Flunarizine effectively inhibits IR-induced astrocyte reactivity *in vitro* without inducing cytotoxicity, in part by suppressing key signaling pathways implicated in astrocyte activation such as TGF-β.

### Flunarizine inhibits IR-induced astrocyte reactivity *in vivo* across dieerent models

To evaluate whether Flunarizine can inhibit IR-induced astrocyte reactivity *in vivo*, we used an RCAS/*tv-a*-based glioma model in which PDGFB and a shRNA targeting Tp53 are expressed in Nestin-positive neural progenitor cells in the neonatal mouse brain (**Fig. 4A**). This model recapitulates the histological and genetic features of human gliomas (46). At 28 days post-injection, half of the animals underwent whole-brain irradiation with a dose of 10 Gy. Flunarizine dihydrochloride or vehicle was administered once daily for 3 days before and 3 days after irradiation. Mice were euthanized 72 hours after IR to investigate the short-term response to IR. As expected, 10 Gy IR dramatically reduced tumor cell density; however, Flunarizine treatment did not alter the proportion of surviving tumor cells (**Fig. 4B, 4D**). Neither IR nor Flunarizine affected blood vessel density (**Supp. Fig. 8A**) or the tumor apoptotic index (**Supp. Fig. 8B**). Importantly, GFAP signal-indicative of astrocyte reactivity-was markedly increased in the tumors that received 10 Gy/vehicle in comparison with 0 Gy/vehicle tumors, and this effect was completely abrogated by Flunarizine treatment, confirming its ability to inhibit the induction of astrocyte reactivity *in vivo* (**Fig. 4C-D**).

**Figure 4:**
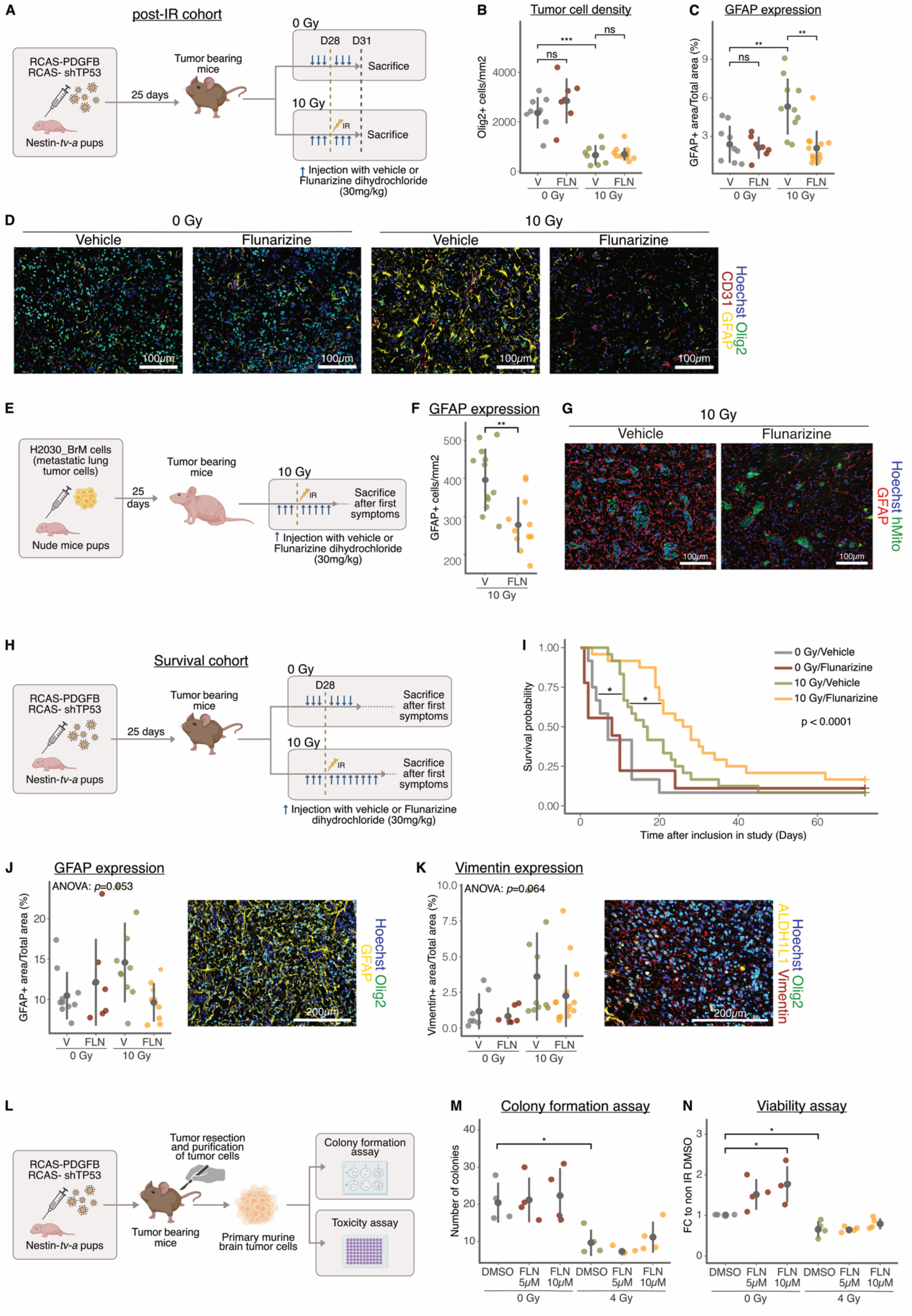
Flunarizine inhibits IR-induced astrocyte reactivity *in vivo* and extends survival in combination with IR in a glioblastoma mouse model. **(A)** Schematic of the study design for the post-IR cohort. Mice were treated with Flunarizine (30mg/kg) (FLN) or vehicle (V), received or not whole-brain irradiation with 10 Gy and were euthanized 3 days after irradiation, or at an equivalent time point for non-irradiated groups. **(B-D)** Fluorescent staining of Hoechst, Olig2, GFAP and CD31 was performed on brain sections from mice in the post-IR cohort described in **(A). (D)** Representative images of the staining and quantification of tumor cell density **(B)** and GFAP expression **(C). (E)** Schematic of the study design for the brain metastasis cohort. Mice were treated with Flunarizine (30mg/kg) or vehicle, received whole brain irradiation with 10 Gy and were euthanized upon onset of tumor symptoms. **(F-G)** Representative images **(G)** and quantification of GFAP expression **(F)** in brain sections from mice in the brain metastasis cohort described in **(E). (H)** Schematic of the study design for the GBM survival cohort. Mice were treated with Flunarizine (30mg/kg) or vehicle, received or not whole brain irradiation with 10 Gy and euthanized upon tumor symptom onset. **(I)** Kaplan-Meier survival curves for mice treated with vehicle (*n*=12), Flunarizine (*n*=9), vehicle/10Gy (*n*=24), or Flunarizine/ 10 Gy (*n*=24). Statistical analysis was performed using Gehan-Breslow (generalized Wilcoxon) test, followed by pairwise comparisons using log-rank tests. **(J)** Representative image and quantification of GFAP expression in the survival cohort described in **(H). (K)** Representative image and quantification of vimentin expression in the survival cohort described in **(H). (L)** Schematic of the purification of primary murine GBM cells used for the colony formation assay **(M)** and toxicity assay **(N). (M)** Colony formation assay using primary murine GBM cells treated with 0 or 4 Gy and DMSO or Flunarizine (*n*=4). **(N)** WST-1 assay using primary murine GBM cells treated with 0 or 4 Gy and DMSO or Flunarizine. Results are expressed as fold change relative to non-IR DMSO control (*n*=4). Data in this figure show mean ± SD. *P*-values were determined using one-way ANOVA followed by post-hoc unpaired *t*-tests if significant. \**P* < 0.05, \*\**P* < 0.01, \*\*\**P* < 0.001. For all Flunarizine treatments in these experiments, Flunarizine dihydrochloride was used.

To further validate these findings, we utilized another *in vivo* model, that forms brain metastasis from metastatic lung tumor cells. We used the metastatic cell line H2030_BrM, which was generated previously by repeated *in vivo* selection of the cells with ability to form brain metastasis after intra-cardiac inoculation (34). When injected intracranially into immunocompromised *Foxn1*_nu/nu_ pups, these cells with high tropism for brain tissue reliably formed brain tumors within a few weeks (**Fig. 4E**). All mice received 10 Gy IR in combination with either vehicle or Flunarizine treatment and were euthanized upon the appearance of tumor symptoms. Although Flunarizine did not improve overall survival (**Supp. Fig 9**), we observed a significant reduction of GFAP+ cells in tumors from Flunarizine-treated mice compared to controls (**Fig. 4F**). These results confirmed that Flunarizine effectively inhibits astrocyte reactivity *in vivo* and its efficacy across two distinct tumor models suggests that its effect is independent of tumor type.

### Flunarizine increases survival in combination with IR in a glioblastoma mouse model

To assess the therapeutic potential of Flunarizine for GBM, we conducted a pre-clinical study using the RCAS/tv-a glioma model described previously. A similar treatment protocol was followed, but in this experiment, mice were monitored until they developed tumor-related symptoms (**Fig. 4H**). Flunarizine alone did not increase survival compared to vehicle control (**Fig. 4I**). As expected, treatment with 10 Gy IR significantly prolonged mouse survival relative to non-irradiated controls. Notably, the combination of Flunarizine and IR further extended survival compared to IR alone.

Histological analysis revealed that neither Flunarizine nor IR altered tumor cell density (**Supp. Fig. 10A-B**) or overall astrocyte density, assessed by the pan-astrocyte marker ALDH1L1 (**Supp. Fig. 10A-C**). Recurrent tumors exhibited a higher Ki67 proliferative index compared to primary tumors, but this was unaffected by Flunarizine (**Supp. Fig. 10A-D**). Similarly, no significant differences were observed across treatment groups in apoptotic index (**Supp. Fig. 10E-F**), necrotic area (**Supp. Fig. 10E-G**) or vascular density (**Supp. Fig. 10H-I**). When investigating the long-term effect of Flunarizine after on astrocyte reactivity, we observed a non-significant increase in GFAP and Vimentin expression in 10 Gy/vehicle-treated tumors, which was partially (for Vimentin) or totally (for GFAP) reversed in 10 Gy/Flunarizine tumors (Fig. 4J-K). These findings suggest that Flunarizine effectively inhibits IR-induced astrocyte reactivity, with a sustained but attenuated effect over time.

To determine whether the survival benefit of Flunarizine in combination with IR could be attributed to a direct effect on tumor cells, we purified primary tumor cells from the RCAS/tv-a glioma model described previously (**Fig. 4L**) and subjected them to colony formation and toxicity assays. Flunarizine, alone or combined with IR, did not reduce colony formation (**Fig. 4M**) or affect tumor cell viability (**Fig. 4N**). We observed similar results in colony formation assays using the GBM cell lines U251 and T98G (**Supp. Fig. 11A-B**) and in toxicity using U251MG cell line (**Supp. Fig. 11C**) as well as serum-free cultured human glioma cell lines U3020MG, U3065MG and U3082MG (**Supp. Fig. 11D-F**).

Together, these results demonstrate that Flunarizine significantly improves survival when combined with IR *in vivo*, likely by targeting the tumor microenvironment and more specifically reactive astrocytes, rather than exerting direct cytotoxic effects on tumor cells.

### IR-induced scar formation is inhibited by Flunarizine

To understand how Flunarizine delays recurrence *in vivo*, we investigated its effect when combined with IR on ECM deposition. A recent study showed that GBM forms a fibrotic scar following treatment, including after RT, which contributes to tumor recurrence (47). We stained tumors from the survival study (**Fig. 4H**) for type I collagen and observed increased collagen I deposition in recurrent tumors. This effect was attenuated by Flunarizine treatment (**Fig. 5A-B**). We found a strong enrichment of GFAP signal in regions covering or surrounding collagen I (**Fig C-D**), suggesting a contribution of reactive astrocytes to fibrotic scar formation.

**Figure 5:**
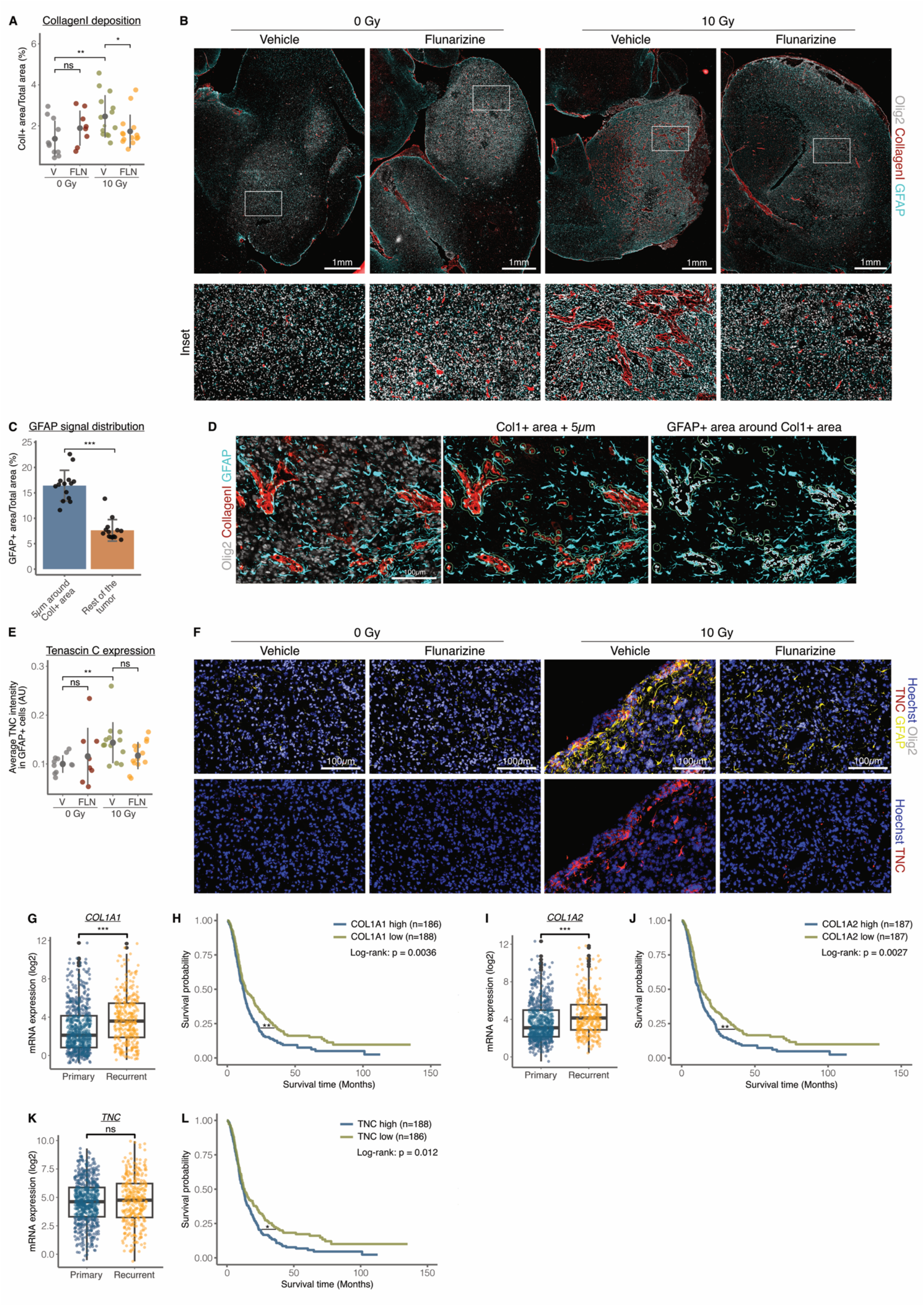
Flunarizine inhibits IR-induced fibrotic scar formation in a mouse model of glioblastoma. **(A-D)** Immunofluorescent staining of Olig2, GFAP and Collagen I on brain sections from mice in the survival cohort (described in Fig. 4H). **(A)** Quantification of collagen I deposition and **(B)** representative images of the staining. **(C)** Quantification of GFAP signal in Vehicle/10 Gy mice, comparing its distribution between peri-Collagen I areas (5 µm around Col1^+^ regions) and tumor bulk (remaining tumor area). Statistical significance was assessed using unpaired t-tests. **(D)** Representative images showing total staining (left), identification of peri-Collagen I areas (middle), and detection of GFAP signal in those areas (right). **(E-F)** Immunofluorescent staining of Olig2, GFAP and TNC performed on brain sections from mice in the survival cohort (described in Figure 4H). **(E)** Quantification of TNC signal in GFAP+ cells and **(F)** representative images of the staining. Fluorescence intensity is shown in arbitrary units (AU). **(G, I, K)** Expression levels of *COL1A1* **(G)**, *COL1A2* **(I)** and *TNC* **(K)** in primary versus recurrent GBM using the data from Chinese Glioma Genome Atlas (CGGA) obtained via the Gliovis portal. **(H, J, L)** Kaplan-Meier survival curves of GBM patients with low versus high *COL1A1* **(H)**, *COL1A2* **(J)** or *TNC* **(L)** expression using the data from Chinese Glioma Genome Atlas (CGGA) obtained using the Gliovis portal. Statistical significance was determined using log-rank tests. **(A, E)** Data are shown as mean ± SD. *P*-values were determined using one-way ANOVA followed by post-hoc unpaired *t*-tests if significant. \**P* < 0.05, \*\**P* < 0.01, \*\*\**P* < 0.001.

RNA-sequencing data showed that adding Flunarizine to IR treatment suppressed the expression of Tenascin C (*TNC*), another ECM protein (**Fig. 3E**). Immunostaining revealed strong TNC expression in GFAP+ astrocytes. Quantification indicated an increase of the TNC signal in GFAP+ cells from 10 Gy/vehicle-treated tumors, which was partially reversed in the 10Gy/Flunarizine group, although the difference did not reach statistical significance (**Fig. 5 E-F**). Notably, there was no significant differences in collagen I (**Supp. Fig. 12A**) or TNC deposition (**Supp. Fig. 12B**), between treatment groups in the post-IR cohort (**Fig. 4A**), suggesting that ECM protein accumulation becomes detectable only after tumor recurrence.

These findings support the idea that Flunarizine inhibits IR-induced fibrotic scar formation, a process that may contribute to tumor recurrence. To assess the clinical relevance of this mechanism, we examined the expression of *COL1A1* and *COL1A2* (coding for collagen type 1) and *TNC* in primary versus recurrent tumors using Gliovis (37) to access bulk RNA-seq datasets from the Chinese Glioma Genome Atlas (CGGA) focusing on GBM patients. Both *COL1A1* and *COL1A2* were significantly overexpressed in patients in recurrent tumors compared to primary tumors (**Fig. 5G, 5I**), and their high expression was associated with shorter patient survival (**Fig. 5H, 5J**). While *TNC* was not overexpressed in recurrent tumors (**Fig. 5K**), its high expression was also associated with poor prognosis (**Fig. 5L**). Furthermore, all three genes were preferentially overexpressed in tumors with wild-type IDH and without 1p/19q codeletion (**Supp. Fig. 13A-C**), supporting their relevance in aggressive glioblastoma subtypes.

Based on our findings, we propose a model in which Flunarizine inhibits IR-induced astrocyte reactivity shortly after IR. This inhibition prevents the formation of the fibrotic scar composed of ECM proteins such as collagen I and TNC, thereby delaying tumor recurrence and extending survival in a preclinical mouse model of GBM (**Fig. 6**).

**Figure 6:**
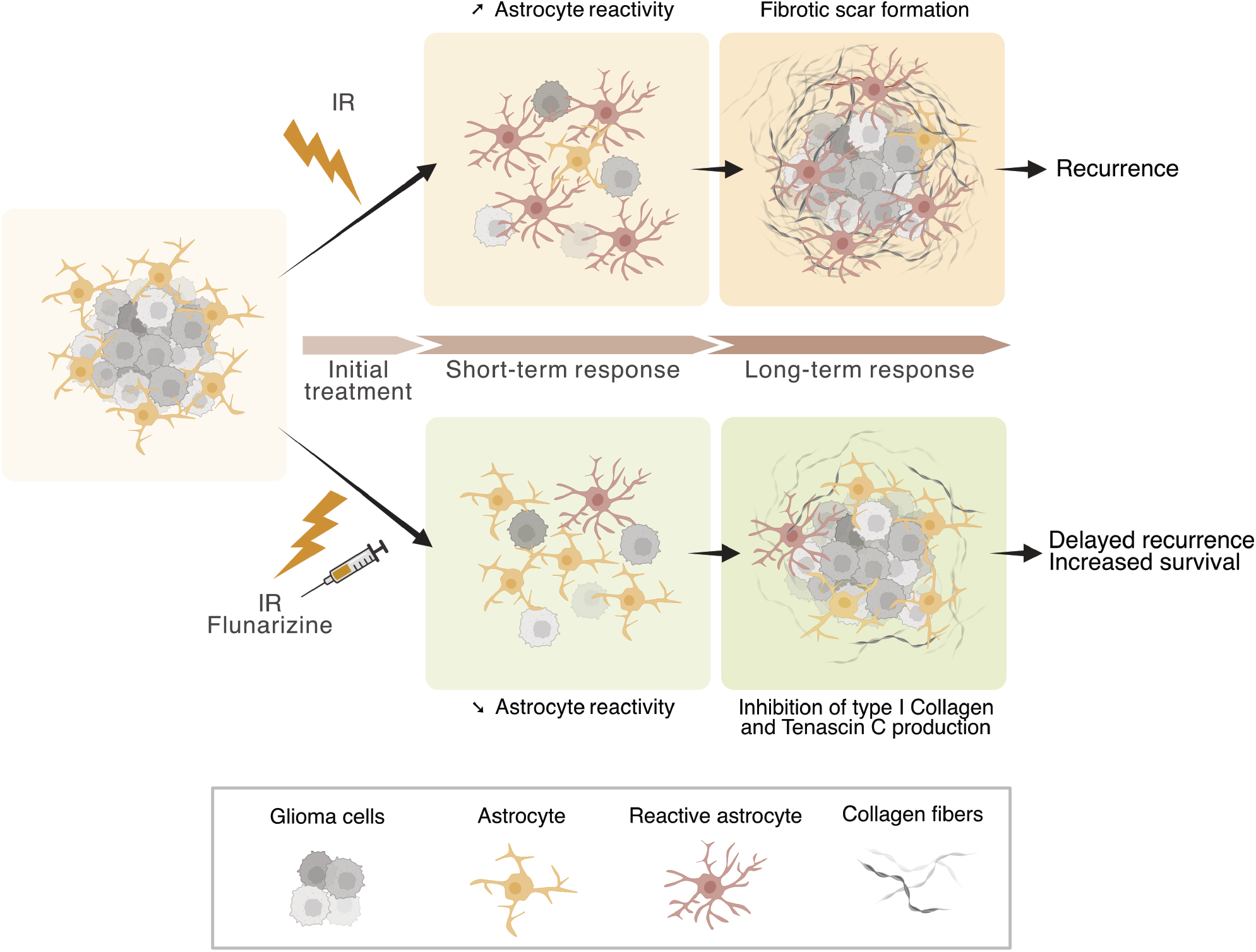
Summary of the findings. Proposed model in which Flunarizine inhibits the early induction of astrocyte reactivity following IR, thereby preventing the formation of a fibrotic scar composed of Collagen I and Tenascin C, which otherwise promotes tumor recurrence.

## Discussion

Astrocyte reactivity is an evolutionarily conserved process that plays a crucial role in protecting brain tissue from a wide range of insults. However, excessive, insufficient, or aberrant activation of astrocytes has been observed in-and, in some cases, directly implicated in-many different types of diseases (48) including neurodegenerative diseases, autoimmune conditions, stroke, traumatic brain injury, and both primary and secondary brain tumors. Identifying compounds that modulate astrocyte reactivity holds significant therapeutic potential across these contexts. Over the past decades, efforts to understand astrocyte function and dysfunction have revealed challenges in identifying consistent markers of reactivity that are shared across disease states (5). The only universal hallmark of astrocyte reactivity observed across most conditions is a profound change in cellular morphology following reactivity-inducing stimuli (5). Building on this, we developed a novel *in vitro* approach to monitor astrocyte reactivity focusing on morphology changes in both the cell body and nucleus, as well as the expression, subcellular localization and signal pattern of two established reactivity markers. We validated this platform using a range of reactivity inducers and inhibitors. While changes in astrocyte morphology have traditionally served as a qualitative and empirical readout of reactivity, our approach provides an automated, unbiased, and robust method for quantification. Notably, a recent large-scale study has demonstrated that high-content morphological profiling is a powerful approach not only for assessing cellular responses but also for inferring mechanisms of action and compound bioactivity, emphasizing its growing value in drug discovery pipelines (49). In this context, our method extends these principles to astrocyte biology, providing a scalable and informative platform for screening modulators of astrocyte reactivity.

In recent years, a growing number of studies have demonstrated that stromal astrocytes play a pro-tumorigenic role in both primary brain tumors and brain metastases. Notably, astrocytes influence tumor progression through a wide variety of mechanisms. Within the brain tumor microenvironment, they have been shown to support tumor metabolism by fueling tumor cells with cholesterol (10) or transferring mitochondria via gap junctions (11); to promote an immunosuppressive environment by inducing T-cell apoptosis (9) or modulating the recruitment and activation of innate immune cells (8, 10, 12); to regulate tumor dormancy in brain metastasis by inducing quiescence in tumor cells (50); and to facilitate recurrence after RT by remodeling the ECM (14). Targeting these pro-tumoral functions could therefore disrupt tumor growth on multiple fronts, potentially enhancing therapeutic efficacy and reducing the likelihood of resistance. A recent phase I clinical trial using the STAT3 inhibitor Silibinin - targeting STAT3+ reactive astrocytes-showed a significant improvement in overall survival in patients with various types of brain metastasis (12), highlighting the therapeutic potential of astrocyte-directed interventions. A phase-II trial is currently underway to evaluate Silibinin in patients with brain metastases from NSCLC or breast cancer. However, there remains a need to identify additional therapies that can selectively target reactive astrocytes and adapt to the various functional states they adopt in different disease contexts. Our work contributes to this effort by providing a list of promising drug repurposing candidates that target astrocytes activated in response to IR. Since RT is a part of the standard of care for both GBM and brain metastases, targeting IR-induced astrocyte reactivity is highly relevant to these conditions. We have already demonstrated the therapeutic potential of one of these candidates - Flunarizine-in a preclinical model, validating its efficacy in improving survival outcomes. Notably, Flunarizine is an off-patent compound, that has been approved for decades to treat migraines and is associated with only mild side effects (41, 42), making it an especially attractive candidate for drug repurposing in GBM.

Following astrocyte activation, the formation of a fibrotic scar has been observed across a wide range of pathological contexts (51). Previously referred to as “glial scar”, it is now understood that although astrocytes contribute significantly to scar formation, the majority of extracellular matrix (ECM) deposition is produced by fibrotic cells—justifying the updated term “fibrotic scar” (48). Nonetheless, several studies have shown that astrocytes themselves can secrete specific ECM components that may directly promote tumor recurrence (14, 50). Our findings showed that IR induces increased type 1 collagen deposition and elevated TNC expression by reactive astrocytes. We found that the genes coding for both proteins are associated with poorer survival outcome in GBM patients. Type 1 collagen is involved in the formation of “oncostreams” - structural features linked to tumor aggressiveness in glioblastoma - (52), and in the organization of the glioma stem cell niche, further supporting its potential role in driving tumor recurrence (53). TNC has also been implicated in various aspects of glioma progression, including treatment resistance (54). Interestingly, while we observed upregulation of *TNC* by astrocytes, *COL1A1* and *COL1A2* expression were not induced by IR in our *in vitro* RNA-seq data. This suggests that astrocytes may contribute to type I collagen deposition indirectly, possibly by stimulating fibrotic stromal cells within the tumor microenvironment.

In conclusion, our study validated the pro-tumorigenic role of astrocyte reactivity in the context of GBM and provided evidence linking the formation of a fibrotic scar to tumor recurrence following RT. We introduced a novel, automated method to monitor astrocyte reactive states *in vitro*, enabling more robust and unbiased assessment of this complex phenotype. Finally, we identified Flunarizine as a repurposable compound that effectively inhibits IR-induced astrocyte reactivity and delays tumor recurrence, offering a promising therapeutic strategy to enhance the efficacy of current GBM treatments.

## Acknowledgements

We thank Katarzyna Radke and Elinn Johansson for valuable input. We acknowledge Clinical Genomics Lund, SciLifeLab and Center for Translational Genomics (CTG), Lund University, for providing expertise and service with sequencing and analysis. We acknowledge the High-Content CRISPR Screens (HCCS) facility in Biotech Research & Innovation Centre (BRIC), Copenhagen, Denmark, for providing expertise and service with drug screening. We thank Joan Massagué (Memorial Sloan Kettering Cancer Center, New York, USA) for kindly sharing with us the H2030_BrM3 cells.

## References

1. Stupp R, et al. Radiotherapy plus Concomitant and Adjuvant Temozolomide for Glioblastoma. New England Journal of Medicine. 2005;352(10):987–996.

2. Silva MID, Stringer BW, Bardy C. Neuronal and tumourigenic boundaries of glioblastoma plasticity. Trends in Cancer. 2023;9(3):223–236.

3. Read RD, et al. Glioblastoma microenvironment—from biology to therapy. Genes Dev. 2024;38(9–10):360–379.

4. Berg TJ, Pietras A. Radiotherapy-induced remodeling of the tumor microenvironment by stromal cells. Seminars in Cancer Biology. 2022;86:846–856.

5. Escartin C, et al. Reactive astrocyte nomenclature, definitions, and future directions. Nat Neurosci. 2021;24(3):312–325.

6. Wu J, et al. Reactive Astrocytes in Glioma: Emerging Opportunities and Challenges. International Journal of Molecular Sciences. 2025;26(7):2907.

7. Parmigiani E, et al. Old Stars and New Players in the Brain Tumor Microenvironment. Front Cell Neurosci. 2021;15. 10.3389/fncel.2021.709917.

8. Henrik Heiland D, et al. Tumor-associated reactive astrocytes aid the evolution of immunosuppressive environment in glioblastoma. Nat Commun. 2019;10(1):2541.

9. Faust Akl C, et al. Glioblastoma-instructed astrocytes suppress tumour-specific T cell immunity. Nature. 2025;1–11.

10. Perelroizen R, et al. Astrocyte immunometabolic regulation of the tumour microenvironment drives glioblastoma pathogenicity. Brain. 2022;145(9):3288–3307.

11. Watson DC, et al. GAP43-dependent mitochondria transfer from astrocytes enhances glioblastoma tumorigenicity. Nat Cancer. 2023;4(5):648–664.

12. Priego N, et al. STAT3 labels a subpopulation of reactive astrocytes required for brain metastasis. Nat Med. 2018;24(7):1024–1035.

13. Qu F, et al. Crosstalk between small-cell lung cancer cells and astrocytes mimics brain development to promote brain metastasis. Nat Cell Biol. 2023;25(10):1506–1519.

14. Berg TJ, et al. The Irradiated Brain Microenvironment Supports Glioma Stemness and Survival via Astrocyte-Derived Transglutaminase 2. Cancer Res. 2021;81(8):2101–2115.

15. Canals I, et al. Rapid and efficient induction of functional astrocytes from human pluripotent stem cells. Nat Methods. 2018;15(9):693–696.

16. Canals I, et al. Astrocyte dysfunction and neuronal network hyperactivity in a CRISPR engineered pluripotent stem cell model of frontotemporal dementia. Brain Commun. 2023;5(3):fcad158.

17. Xie Y, et al. The Human Glioblastoma Cell Culture Resource: Validated Cell Models Representing All Molecular Subtypes. eBioMedicine. 2015;2(10):1351–1363.

18. Pietras A, et al. Osteopontin-CD44 Signaling in the Glioma Perivascular Niche Enhances Cancer Stem Cell Phenotypes and Promotes Aggressive Tumor Growth. Cell Stem Cell. 2014;14(3):357–369.

19. Stirling DR, et al. CellProfiler 4: improvements in speed, utility and usability. BMC Bioinformatics. 2021;22(1):433.

20. Lord SJ, et al. SuperPlots: Communicating reproducibility and variability in cell biology. Journal of Cell Biology. 2020;219(6):e202001064.

21. Stirling DR, Carpenter AE, Cimini BA. CellProfiler Analyst 3.0: accessible data exploration and machine learning for image analysis. Bioinformatics. 2021;37(21):3992–3994.

22. Caicedo JC, et al. Data-analysis strategies for image-based cell profiling. Nat Methods. 2017;14(9):849–863.

23. Omta WA, et al. HC StratoMineR: A Web-Based Tool for the Rapid Analysis of High-Content Datasets. Assay Drug Dev Technol. 2016;14(8):439–452.

24. Ewels PA, et al. The nf-core framework for community-curated bioinformatics pipelines. Nat Biotechnol. 2020;38(3):276–278.

25. Patro R, et al. Salmon provides fast and bias-aware quantification of transcript expression. Nat Methods. 2017;14(4):417–419.

26. Soneson C, Love MI, Robinson MD. Differential analyses for RNA-seq: transcript-level estimates improve gene-level inferences [preprint]. 2016. 10.12688/f1000research.7563.2.

27. Love MI, Huber W, Anders S. Moderated estimation of fold change and dispersion for RNA-seq data with DESeq2. Genome Biology. 2014;15(12):550.

28. Durinck S, et al. Mapping identifiers for the integration of genomic datasets with the R/Bioconductor package biomaRt. Nat Protoc. 2009;4(8):1184–1191.

29. Liberzon A, et al. The Molecular Signatures Database Hallmark Gene Set Collection. Cell Systems. 2015;1(6):417–425.

30. Badia-i-Mompel P, et al. decoupleR: ensemble of computational methods to infer biological activities from omics data. Bioinformatics Advances. 2022;2(1):vbac016.

31. Schubert M, et al. Perturbation-response genes reveal signaling footprints in cancer gene expression. Nat Commun. 2018;9(1):20.

32. Holland EC, et al. A constitutively active epidermal growth factor receptor cooperates with disruption of G1 cell-cycle arrest pathways to induce glioma-like lesions in mice. Genes Dev. 1998;12(23):3675–3685.

33. Kassambara A, et al. survminer: Drawing Survival Curves using “ggplot2” R package version 0.5.0 [Internet]. https://rpkgs.datanovia.com/survminer/index.html. Accessed July 3, 2025.

34. Nguyen DX, et al. WNT/TCF Signaling through LEF1 and HOXB9 Mediates Lung Adenocarcinoma Metastasis. Cell. 2009;138(1):51–62.

35. Bankhead P, et al. QuPath: Open source software for digital pathology image analysis. Sci Rep. 2017;7(1):16878.

36. Zhao Z, et al. Comprehensive RNA-seq transcriptomic profiling in the malignant progression of gliomas. Sci Data. 2017;4:170024.

37. Bowman RL, et al. GlioVis data portal for visualization and analysis of brain tumor expression datasets. Neuro-Oncology. 2017;19(1):139–141.

38. Pantazopoulou V, et al. Hypoxia-Induced Reactivity of Tumor-Associated Astrocytes Affects Glioma Cell Properties. Cells. 2021;10(3):613.

39. Povirk LF. DNA damage and mutagenesis by radiomimetic DNA-cleaving agents: bleomycin, neocarzinostatin and other enediynes. Mutation Research/Fundamental and Molecular Mechanisms of Mutagenesis. 1996;355(1):71–89.

40. Chambers JW, et al. Small Molecule c-jun-N-Terminal Kinase Inhibitors Protect Dopaminergic Neurons in a Model of Parkinson’s Disease. ACS Chem Neurosci. 2011;2(4):198–206.

41. Stubberud A, et al. Flunarizine as prophylaxis for episodic migraine: a systematic review with meta-analysis. PAIN. 2019;160(4):762.

42. Holmes B, et al. Flunarizine. Drugs. 1984;27(1):6–44.

43. Liu B, et al. Epidermal Growth Factor Receptor Activation: An Upstream Signal for Transition of Quiescent Astrocytes into Reactive Astrocytes after Neural Injury. J Neurosci. 2006;26(28):7532–7540.

44. Li Z-W, et al. Inhibiting epidermal growth factor receptor attenuates reactive astrogliosis and improves functional outcome after spinal cord injury in rats. Neurochemistry International. 2011;58(7):812–819.

45. Luo J. TGF-β as a Key Modulator of Astrocyte Reactivity: Disease Relevance and Therapeutic Implications. Biomedicines. 2022;10(5):1206.

46. Dai C, et al. PDGF autocrine stimulation dedifferentiates cultured astrocytes and induces oligodendrogliomas and oligoastrocytomas from neural progenitors and astrocytes in vivo. Genes Dev. 2001;15(15):1913–1925.

47. Watson SS, et al. Fibrotic response to anti-CSF-1R therapy potentiates glioblastoma recurrence. Cancer Cell. 2024;42(9):1507–1527.e11.

48. Verkhratsky A, et al. Astrocytes in human central nervous system diseases: a frontier for new therapies. Sig Transduct Target Ther. 2023;8(1):1–37.

49. Wolff C, et al. Morphological profiling data resource enables prediction of chemical compound properties. iScience. 2025;28(5). 10.1016/j.isci.2025.112445.

50. Dai J, et al. Astrocytic laminin-211 drives disseminated breast tumor cell dormancy in brain. Nat Cancer. 2022;3(1):25–42.

51. Sofroniew MV, Vinters HV. Astrocytes: biology and pathology. Acta Neuropathol. 2010;119(1):7–35.

52. Comba A, et al. Spatiotemporal analysis of glioma heterogeneity reveals COL1A1 as an actionable target to disrupt tumor progression. Nat Commun. 2022;13(1):3606.

53. Motegi H, et al. Type 1 collagen as a potential niche component for CD133-positive glioblastoma cells. Neuropathology. 2014;34(4):378–385.

54. Fu Z, et al. Matricellular protein tenascin C: Implications in glioma progression, gliomagenesis, and treatment. Front Oncol. 2022;12. 10.3389/fonc.2022.971462.

