## Supplementary Figures for "Phenotypic Screening Identifies Flunarizine as an Inhibitor of Radiotherapy-Induced Astrocyte Reactivity with Therapeutic Potential in Glioblastoma"

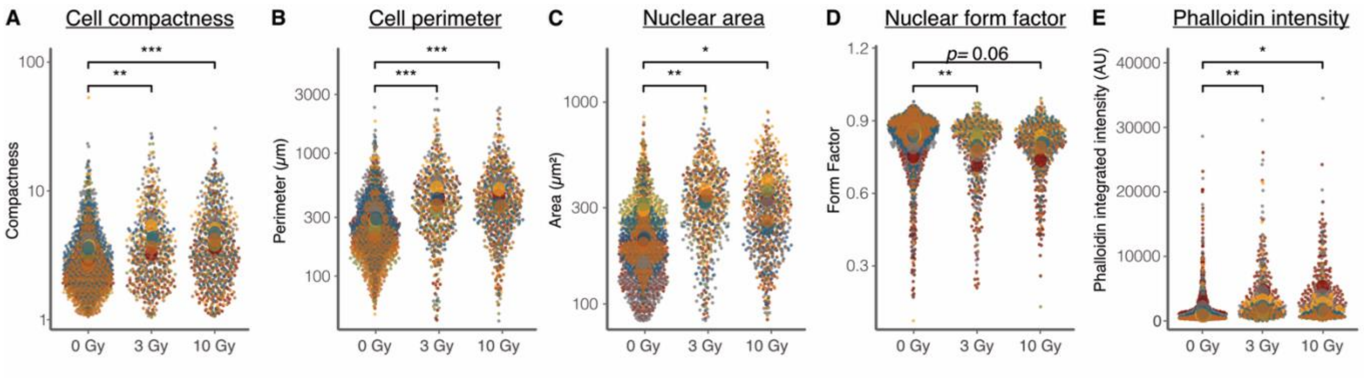


**Supplementary Figure 1: (A-E)** Human astrocytes were irradiated 0, 3 or 10 Gy and stained for DNA (Hoechst-33342), F-Actin (Phalloidin), vimentin and GFAP. Acquired images were used to quantify Cell compactness **(A)**, cell perimeter **(B)**, nuclear area **(C)**, nuclear form factor **(D)** and phalloidin intensity **(E)** (*n*=3). Results are presented as SuperPlots, with small dots representing individual measurements and large dots representing means of biological replicate. Each color indicates a distinct biological replicate. Statistical significance was calculated using paired *t*-tests. **P* < 0.05, ***P* < 0.01, ****P* < 0.001.

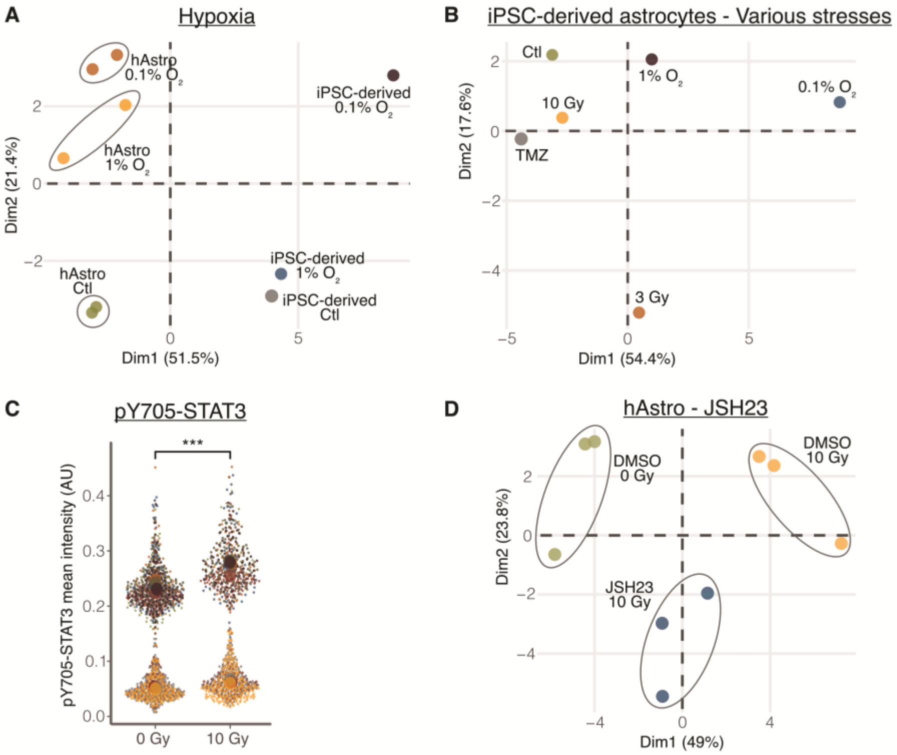


**Supplementary Figure 2: (A, B, D)** PCA of morphological and intensity features extracted as described in Figure 1E comparing: **(A)** response to hypoxia (1% $O_2$ and 0.1% $O_2$) of primary human astrocytes (hAstro) versus iPSC-derived astrocytes (iPSC-derived); **(B)** response of iPSC-derived astrocytes to various stresses vs. control (Ctl) (0 Gy, 21% $O_2$); **(D)** response of primary human astrocytes to 10 Gy and NF-κBi JSH23 (20µM)**.** Experiments analyzed by PCA were repeated at least three times; one representative result is shown. **(C)** Quantification of nuclear pY05-STAT3 mean intensity following immunofluorescent detection in primary human astrocytes treated with 0 or 10 Gy. Results are represented as a SuperPlot, with small dots representing individual measurements and large dots indicating means per biological replicate (*n*=9). Each color indicates a distinct biological replicate. Statistical significance was calculated using paired *t*-tests. ****P* < 0.001

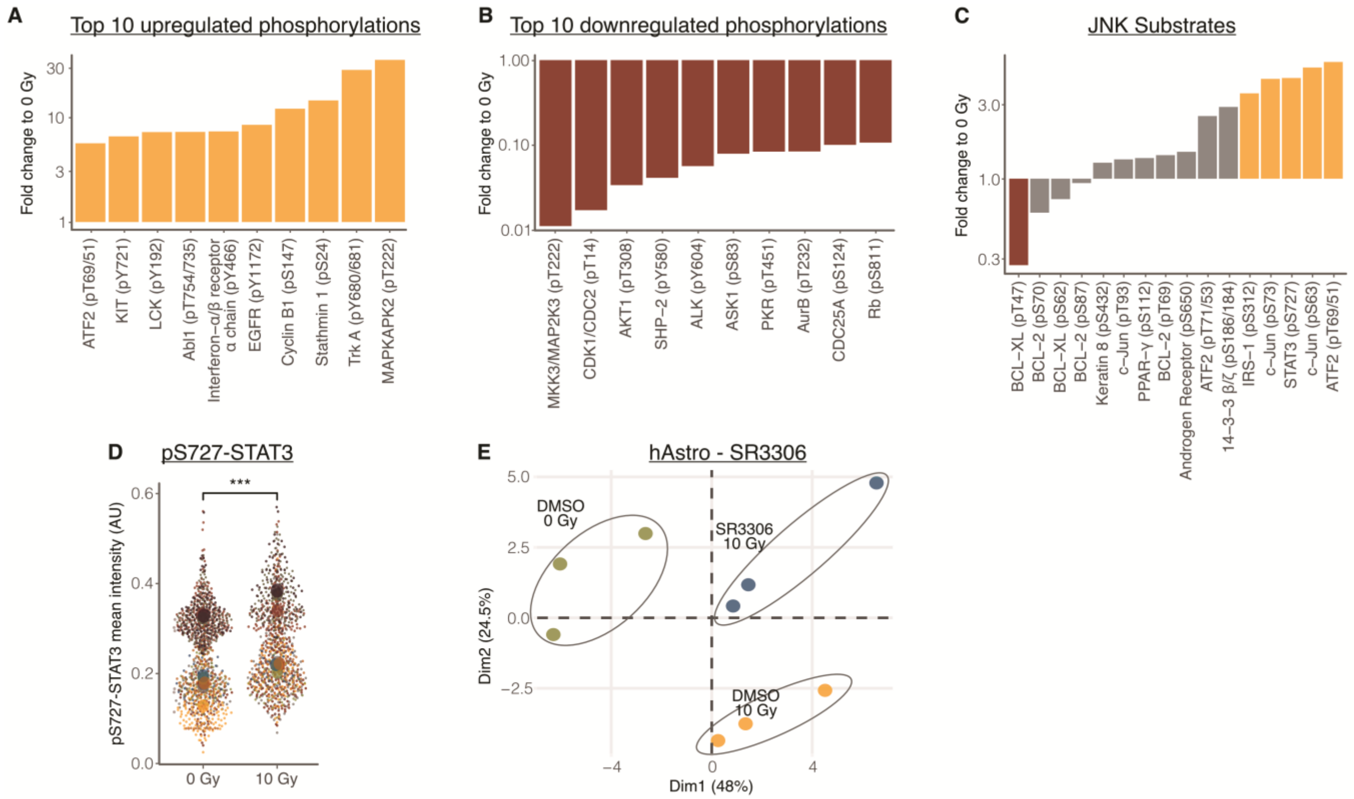


**Supplementary Figure 3: (A-C)** Phospho-protein array performed on primary human astrocytes treated with 10 Gy IR (described in Figure 1I). Results are shown as fold change relative to non-irradiated astrocytes. Bars are colored red if fold change <3, yellow if >3, and gray otherwise. **(A)** Top 10 up-phosphorylated phosphosites. **(B)** Top 10 down-phosphorylated phosphosites. **(C)** All the phosphosites targeted by JNK. **(D)** Quantification of nuclear pS727-STAT3 mean intensity following immunofluorescent detection in primary human astrocytes 24h after receiving 0 or 10 Gy. Results are represented as a SuperPlot, with small dots representing individual measurements and large dots indicating means per biological replicate. Each color indicates a distinct biological replicate (*n*=9). Statistical significance was calculated using paired *t* tests. ****P* < 0.001 **(E)** PCA of the morphological and intensity features extracted as described in Figure 1E comparing response of primary human astrocytes to 10 Gy and JNKi SR3306 (0.5μM) **(D).** Experiments analyzed by PCA were repeated at least three times; one representative result is shown.

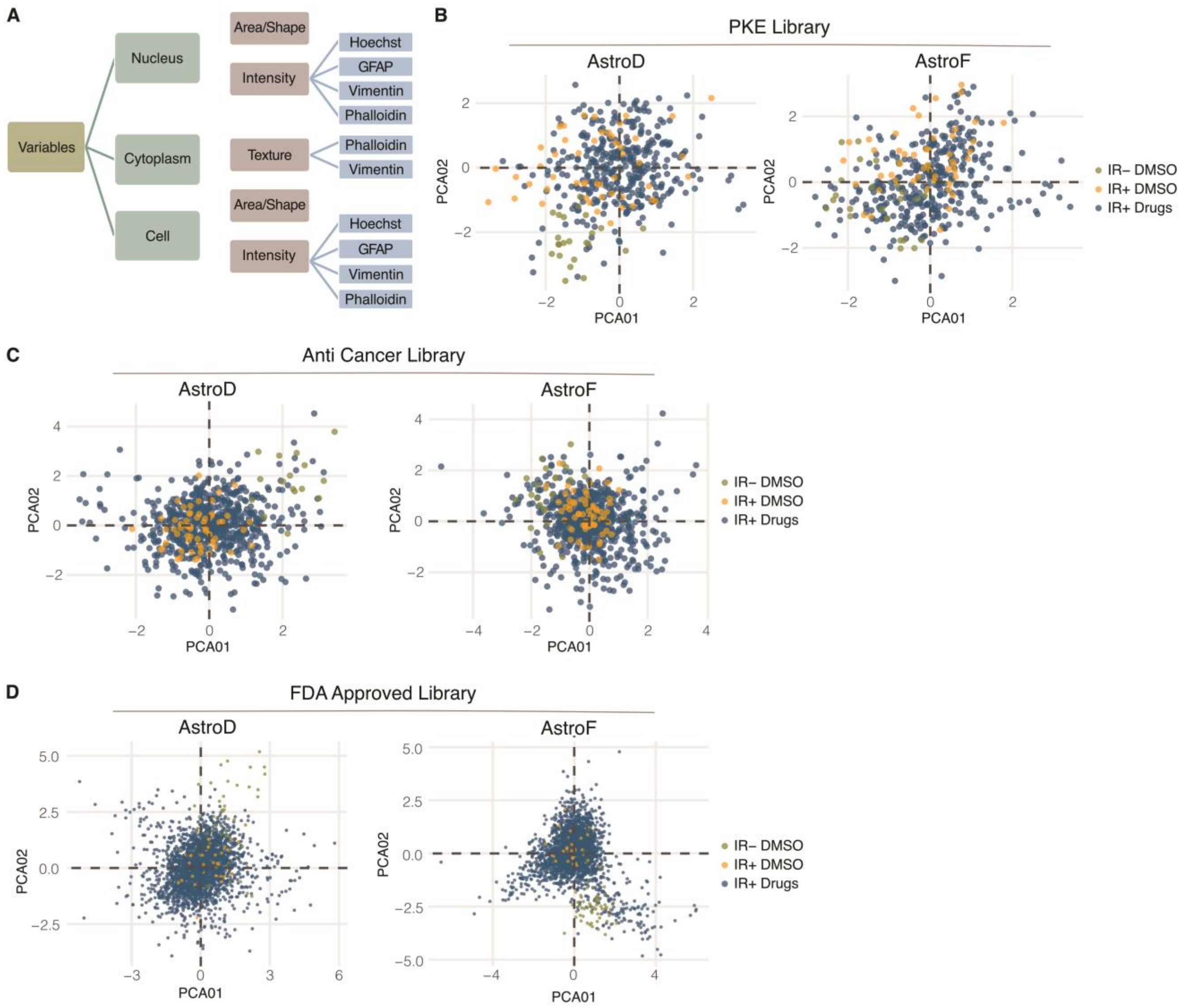


**Supplementary Figure 4: (A)** Schematic illustration of the different types of variables used to perform PCA for the drug screens. **(B-D)** PCA of the morphological and intensity features described in **(A)** comparing response of primary human astrocytes to IR (10 Gy) and compounds from the PKE drug library **(B)**, Anti-Cancer TargetMol library **(C)** and FDA Approved compounds library (Prestwick Chemicals) **(D)** in two different astrocytes batches.

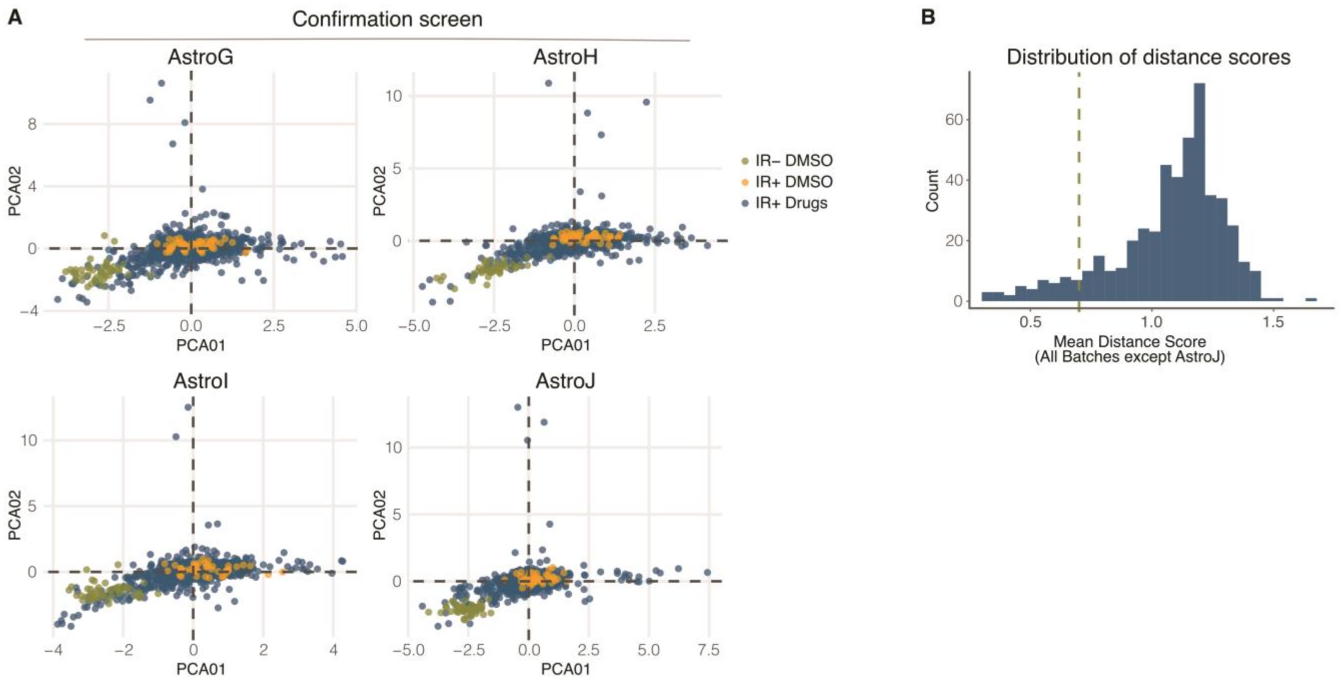


**Supplementary Figure 5: (A)** PCA of the morphological and intensity features described in Supplementary Figure 4A comparing response of primary human astrocytes to IR (10 Gy) and compounds from the confirmation screen in four different astrocytes batches. **(B)** Distribution of the distance scores for the compounds included in the confirmation screen. Each compound was tested in duplicate at four different concentrations. For each concentration, the mean distance score was calculated across three astrocyte batches (AstroG, AstroH, and AstroI) and is shown in the graph.

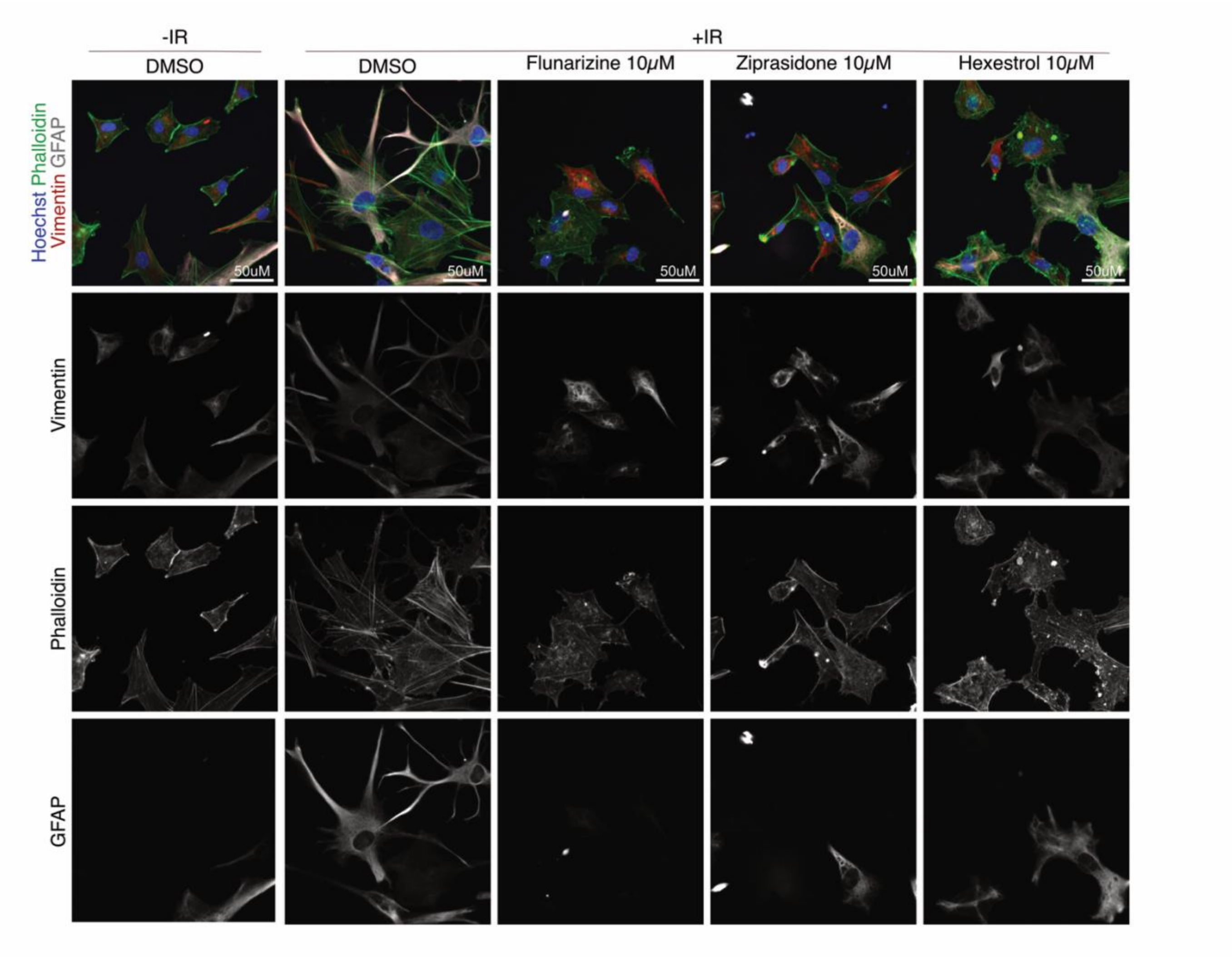


**Supplementary Figure 6:** Representative images from the confirmation screen showing staining of DNA (Hoechst-33342), F-Actin (Phalloidin), vimentin, and GFAP in primary human astrocytes. Astrocytes were treated with 0 or 10 Gy and exposed to selected hit compounds: Flunarizine 10μM, Ziprasidone 10μM Hexestrol 10μM.

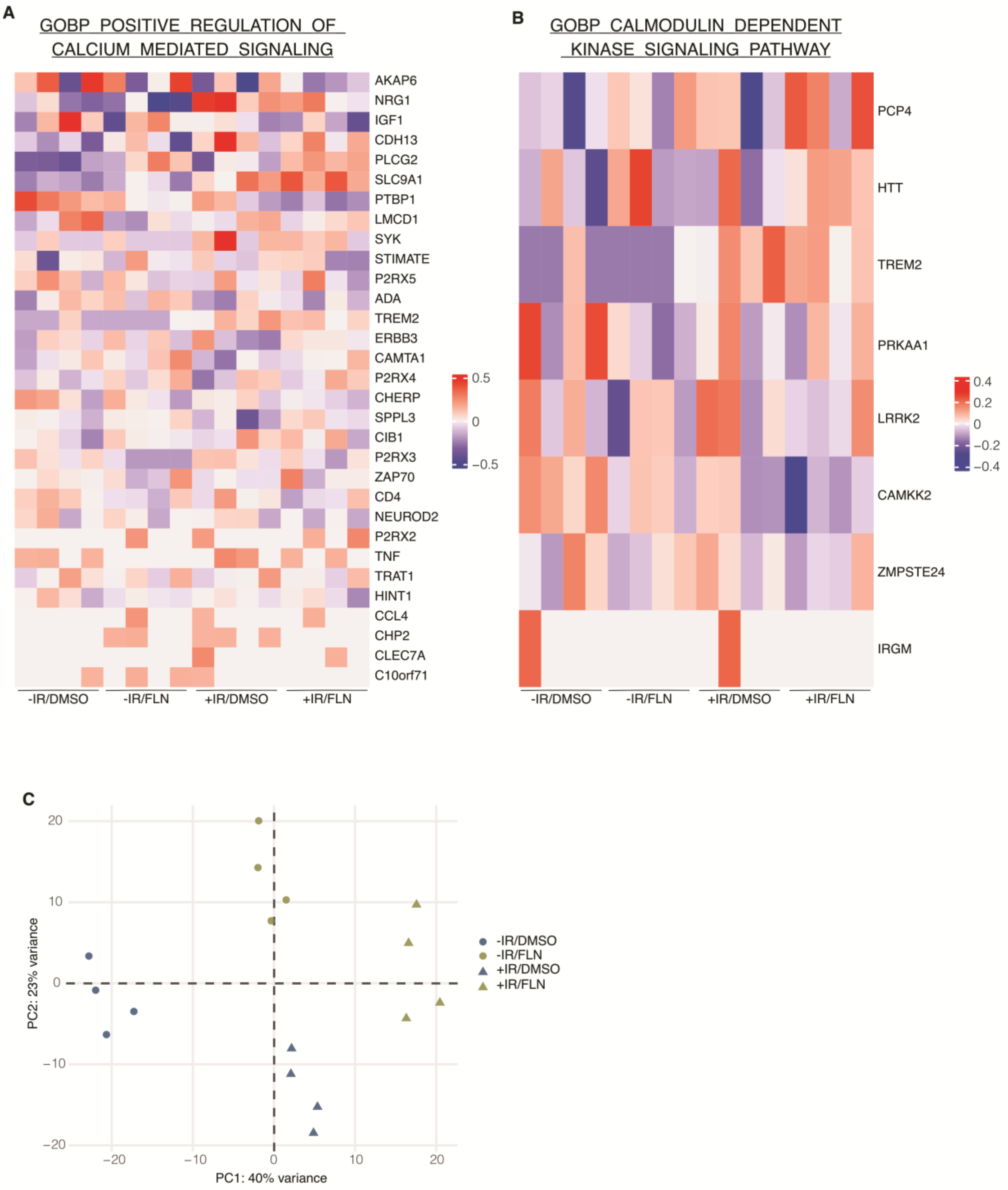


**Supplementary Figure 7: (A-B)** Row-normalized heatmaps of genes from selected Gene Ontology (GO) signatures "positive regulation of calcium-mediated signaling" **(A)** and "calmodulin-dependent kinase signaling" **(B)** across all 16 samples. Data shown as VST-transformed expression. Each row is median-centered and ordered by variance. Expression values are shown as z-scores. (**C**) PCA of VST-transformed expression data (top 5000 most variable genes). Samples colored by treatment and shaped by irradiation condition

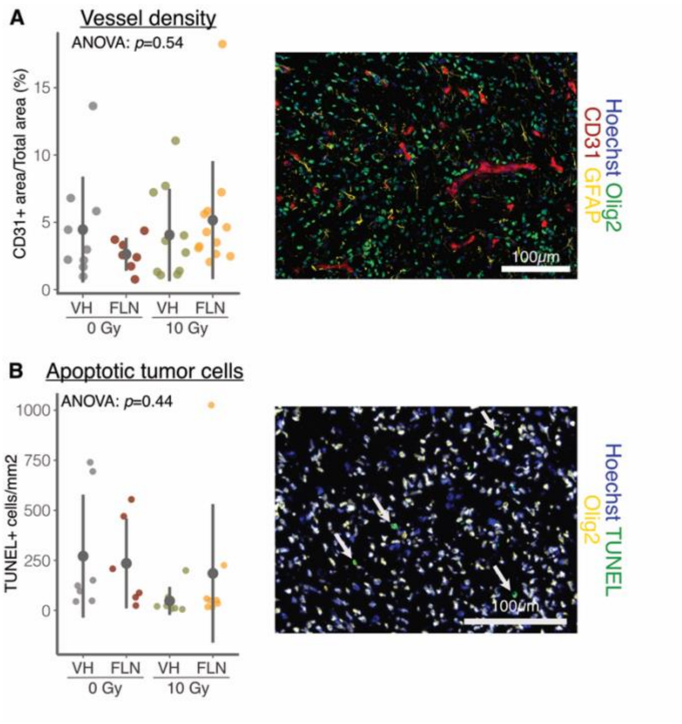


**Supplementary Figure 8: (A-B)** Fluorescent staining on brain sections from the post-IR cohort (described in Figure 4A). **(A)** Quantification of vessel density (left) and representative image showing staining with Hoechst, Olig2, GFAP and CD31. **(B)** Quantification of apoptotic tumor cells (left) and representative image showing staining with Hoechst, Olig2 and TUNEL. Data in this figure show mean ± SD. *P*-values were determined using one-way ANOVA.

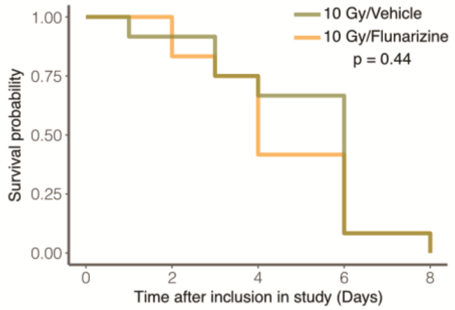


**Supplementary Figure 9:** Kaplan-Meier survival curve for mice treated with vehicle/10Gy ($n$=12), or Flunarizine/10 Gy ($n$=12) in brain metastasis model (described in Figure 4E). Statistical analysis was performed log-rank test.

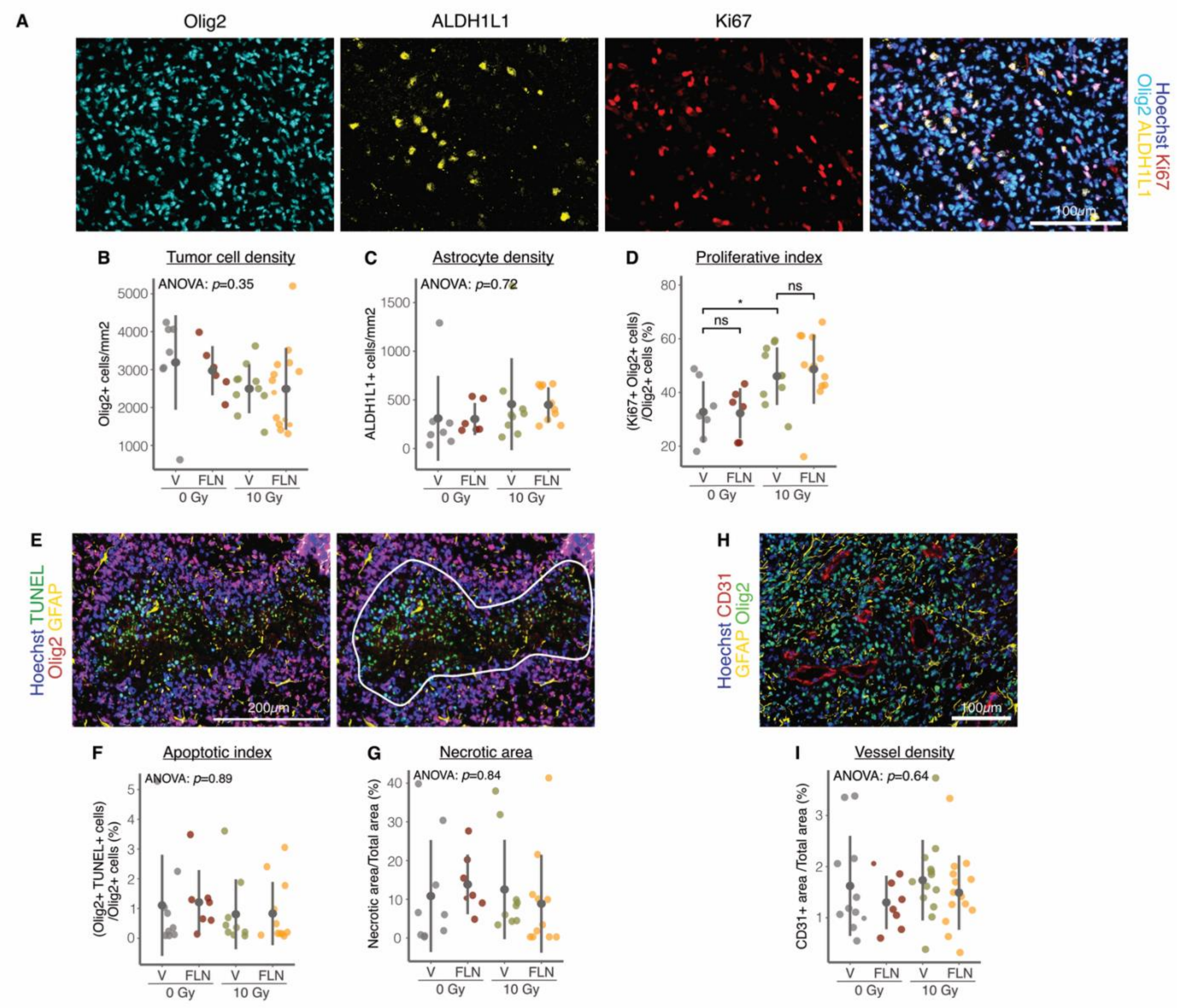


**Supplementary Figure 10: (A-I)** Fluorescent staining on brain sections from the survival cohort (described in Figure 4H). **(A)** Representative images showing staining with Hoechst, Olig2, ALDH1L1 and Ki67. Quantification of tumor cell density **(B)**, astrocyte density **(C)** and tumor proliferative index **(D)**. **(E)** Representative images showing staining with Hoechst, TUNEL, Olig2 and GFAP. Quantification of tumor cells apoptotic index **(F)** and necrotic area **(G)**. **(H)** Representative images showing staining with Hoechst, Olig2, GFAP and CD31. **(I)** Quantification of vessel density. Data in this figure show mean ± SD. *P*-values were determined using one-way ANOVA followed by post-hoc unpaired *t*-tests if significant. *$P < 0.05$.

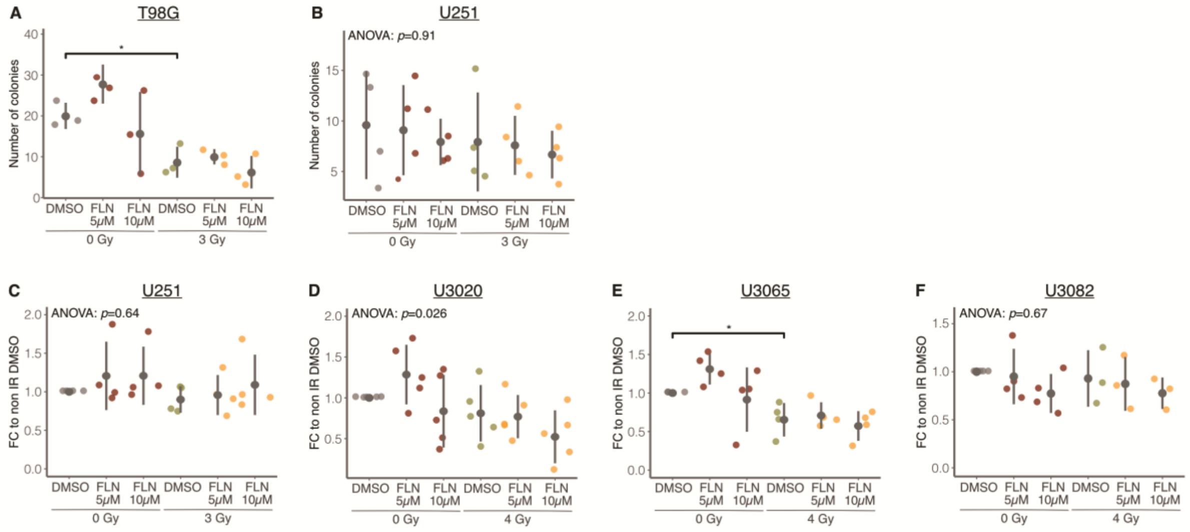


**Supplementary Figure 11: (A-B)** Colony formation assay using T98G (*n*=3) **(A)** or U251 (*n*=4) **(B)** cells treated with 0 or 3 Gy and DMSO or Flunarizine. **(C-F)** WST-1 assay using U251 (*n*=4) **(C)**, U3020 (*n*=5) **(D)**, U3065 (*n*=4) **(E)** or U3082 (*n*=3) **(F)** cells treated with irradiation at indicated dose and DMSO or Flunarizine. Results are expressed as fold change relative to non-IR DMSO control. Data in this figure show mean ± SD. *P*-values were determined using one-way ANOVA followed by post-hoc unpaired *t*-tests if significant. *$P < 0.05$. For all Flunarizine treatments in these experiments, Flunarizine dihydrochloride was used.

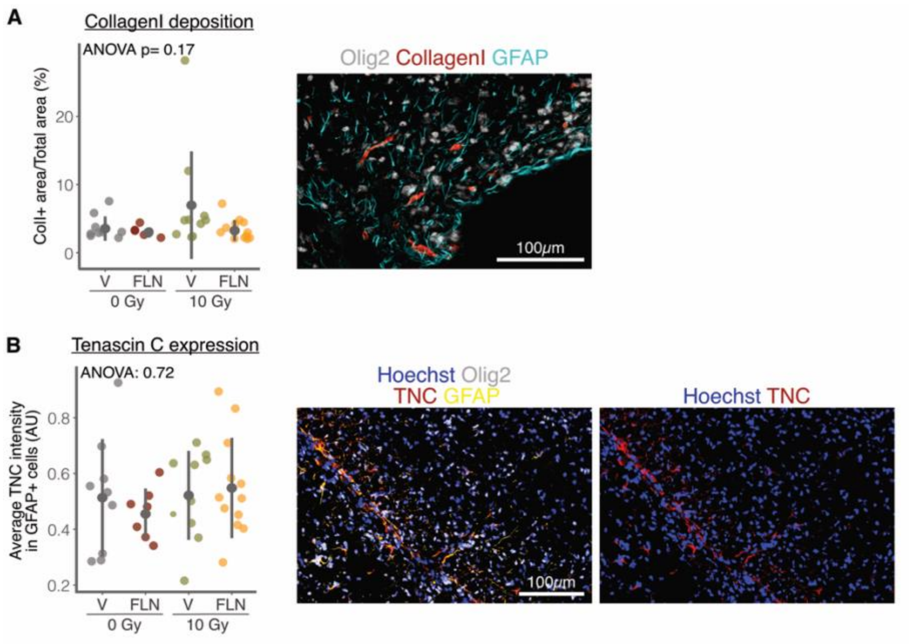


**Supplementary Figure 12: (A-B)** Fluorescent staining on brain sections from the post-IR cohort (described in Figure 4A). **(A)** Quantification of Collagen I deposition (left) and representative image showing staining with Olig2, GFAP and Collagen I. **(B)** Quantification of TNC signal in GFAP+ astrocytes (left) and representative image showing staining with Hoechst, Olig2, GFAP and TNC. Data in this figure show mean ± SD. *P*-values were determined using one-way ANOVA.

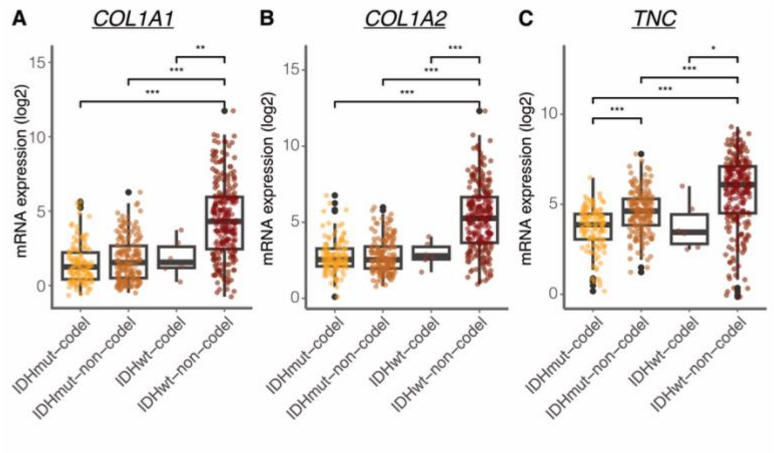


**Supplementary Figure 13: (A-C)** Expression levels of *COL1A1* **(A)**, *COL1A2* **(B)** and *TNC* **(C)** in IDH-WT (IDHwt) glioma compared with IDH mutant (IDHmut) with or without 1p/19q codeletion using the data from Chinese Glioma Genome Atlas (CGGA) obtained via the Gliovis portal. *P*-values were determined using one-way ANOVA followed by post-hoc unpaired *t*-tests if significant. **P* < 0.05 , ***P* < 0.01, ****P* < 0.001.
